# Hi-C guided assemblies reveal conserved regulatory topologies on X and autosomes despite extensive genome shuffling

**DOI:** 10.1101/580969

**Authors:** Gina Renschler, Gautier Richard, Claudia Isabelle Keller Valsecchi, Sarah Toscano, Laura Arrigoni, Fidel Ramirez, Asifa Akhtar

## Abstract

Genome rearrangements that occur during evolution impose major challenges on regulatory mechanisms that rely on three-dimensional genome architecture. Here, we developed a scaffolding algorithm and generated chromosome-length assemblies from Hi-C data for studying genome topology in three distantly related *Drosophila* species. We observe extensive genome shuffling between these species with one synteny breakpoint after approximately every six genes. A/B compartments, a set of large gene-dense topologically associating domains (TADs) and spatial contacts between high-affinity sites (HAS) located on the X chromosome are maintained over 40 million years, indicating architectural conservation at various hierarchies. Evolutionary conserved genes cluster in the vicinity of HAS, while HAS locations appear evolutionarily flexible, thus uncoupling functional requirement of dosage compensation from individual positions on the linear X chromosome. Therefore, 3D architecture is preserved even in scenarios of thousands of rearrangements highlighting its relevance for essential processes such as dosage compensation of the X chromosome.

## Introduction

The development of chromosome conformation capture techniques such as Hi-C, which maps contacts between loci within chromosomes, has facilitated the study of the three-dimensional (3D) architecture of genomes and provided insights into the importance of genome folding for gene regulation (Lieberman-Aiden et al., 2009). Such techniques revealed the existence of genomic regions that show preferential contacts referred to as topologically associating domains (TADs) (Dixon et al., 2012; Nora et al., 2012). TADs and their boundaries also appear to be associated with genomic rearrangements occurring during evolution. Inter-species comparisons in mammals revealed few selected examples, where contiguous orthologous genes inserted at different genomic positions in a different species maintain TAD integrity (Vietri Rudan et al., 2015). This suggests that TADs as modular units are maintained during evolution despite genomic rearrangements. TADs have also been described in non-mammalian species, e.g. *Drosophila* (Sexton et al., 2012), whose genome is more than 10-times smaller, more gene-dense and exposed to faster rates of molecular evolution compared to mammals (Thomas et al., 2010). Due to this reason, *Drosophila* genomes facilitate comparisons between the 3D architecture of highly rearranged, yet related, genomes within a given genus.

Concerted efforts allowed to assemble *Drosophila* genomes (Drosophila 12 Genomes Consortium et al., 2007; Wiegmann and Richards, 2018), however, these assemblies are typically composed of thousands of scaffolds. While such fragmented assemblies are a valuable resource for evolutionary analyses between species, the lack of chromosome-length information hinders comparisons related to genome organization. This, however, would be relevant for the genome-wide analysis of TADs as well as variation of the 3D architecture caused by genomic rearrangements. Recently, information of genome-wide contacts obtained from Hi-C data was utilized for genome assembly leading to the development of several Hi-C scaffolding algorithms. Integrating genome sequencing data with Hi-C-derived information can produce genomes of very high quality and continuity resulting in chromosome-length assemblies. Examples of such published genomes are the mosquito *Aedes aegypti*, the domestic goat *Capra hircus* or the barley *Hordeum vulgare L*. (Bickhart et al., 2017; Burton et al., 2013; Dudchenko et al., 2017; Korbel and Lee, 2013; Marie-Nelly et al., 2014; Mascher et al., 2017).

The observation of genomic rearrangements throughout evolution can raise the question of how they impact mechanisms of transcriptional regulation at genome-wide or chromosome-wide scales. An example of such a chromosome-wide mechanism is dosage compensation, which balances the transcriptional output from sex chromosomes between males and females. In Drosophila, this is achieved by the Male-specific lethal (MSL) complex, which mediates histone H4 lysine 16 acetylation (H4K16ac) resulting in approximately two-fold upregulation of X-linked genes in males (Kuroda et al., 2016; Samata and Akhtar, 2018). This process has been extensively studied in *D. melanogaster* and appears to be conserved in other drosophilids, for example *D. virilis* and *D. busckii*, which are overall separated by approximately 40 million years of evolution (Alekseyenko et al., 2013; Quinn et al., 2016; Robe et al., 2010; Russo et al., 1995). In *D. melanogaster*, the X chromosome adopts a specialized 3D architecture to achieve dosage compensation (Ramírez et al., 2015; Schauer et al., 2017). The X-linked recruitment sites for the MSL complex, termed high-affinity sites (HAS), are enriched in Hi-C contacts between pairs and therefore, appear to cluster together in space.

Here, we studied the evolutionary impact of genomic rearrangements on chromosome conformation by generating Hi-C data in three *Drosophila* species: *D. busckii*, *D. virilis* and *D. melanogaster*. We produced chromosome-length genome assemblies of *D. busckii* and *D. virilis* using Hi-C scaffolding of hybrid contigs assembled *de novo* from PacBio and Illumina reads. To achieve this we developed HiCAssembler, a Hi-C scaffolding tool allowing the assembly of genomes using Hi-C data combined with scaffolds obtained from short and long read sequencing. Using these data and tools, we find extensive rearrangements within chromosomes, while A/B compartments and a subset of TADs appear to be maintained as conserved units. We characterize the set of conserved TADs as being large, dense in genes with essential functions, as well as enriched in histone modifications associated with active transcription. Their boundaries show conserved DNA motifs. Underscoring the functional relevance of maintaining genome topology, we find that spatial contacts implicated in X chromosome dosage compensation are preserved over millions of years of evolution and suggest that they are not a mere consequence of closeness to TAD boundaries or the expression level of their associated genes. Furthermore, we show that genes in the vicinity of HAS are significantly more conserved than genes overlapping with HAS suggesting that the absolute positioning of individual HAS is less important than clustering of genes that require dosage compensation around HAS. Our study in these highly rearranged genomes highlights the importance of maintaining genome topology during evolution, which may shape even chromosome-wide regulatory mechanisms such as on the X chromosome.

## Results

### Chromosome-length assemblies of the *D. busckii* and *D. virilis* genomes

We generated *in situ* Hi-C data from *D. busckii* and *D. virilis* mixed-sex embryos at stage 15-16 to obtain chromosome-length assemblies using Hi-C scaffolding (Figure 1). For *D. busckii*, we additionally generated 2.7 Gb (∼20X genome coverage) PacBio reads of gDNA (Figure S1). We also integrated previously published short Illumina reads (Vicoso and Bachtrog, 2015; Zhou and Bachtrog, 2015) that we assembled into 32,010 short-read contigs using SparseAssembler (Ye et al., 2012). They were then combined with our error-corrected PacBio read data using DBG2OLC (Ye et al., 2016) to obtain a total of 245 contigs with an N50 of 1.4 Mb (Table 1). For *D. virilis,* we used sequence information from the Drosophila 12 Genomes Consortium (Drosophila 12 Genomes Consortium et al., 2007), which contains 13,415 scaffolds (Dvir_caf1, N50 = 10.2 Mb).

**Figure 1.**
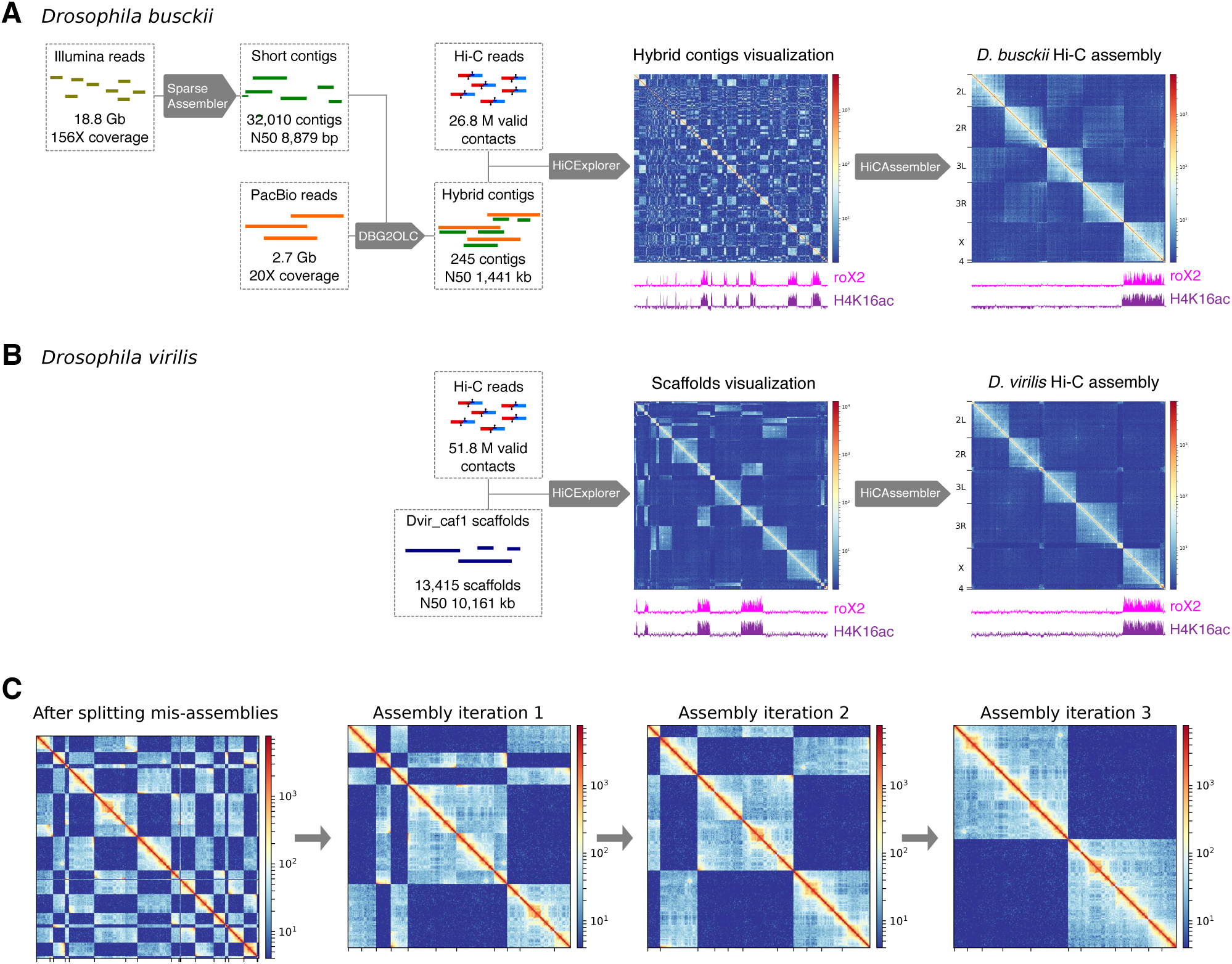
Hi-C guided chromosome-length assemblies of *D. busckii* and *D. virilise* genomes. **(A)** *De novo* assembly of *D. busckii* genome. A hybrid approach integrating long PacBio reads and short contigs assembled from Illumina reads was used to obtain 245 *de novo* contigs of the *D. busckii* genome. Assembly of 156X Illumina reads using SparseAssembler (Ye et al., 2012) resulted in 32,010 short contigs. 20X PacBio data was integrated using DBG2OLC (Ye et al., 2016) which increased the N50 more than 100-fold. These 245 hybrid contigs were scaffolded into chromosome-length with Hi-C data using HiCAssembler. Integrity of the X chromosome (identified by whole genome alignment to *D. melanogaster*) was validated using ChIRP-seq data of the dosage compensation complex member roX2 (Quinn et al., 2016) and ChIP-seq data of H4K16ac from male *D. busckii* larvae. **(B)** *D. virilis* Hi-C assembly. The existing reference scaffolds of *D. virilis* (Dvir_caf1 scaffolds) were assembled into full chromosomes using HiCAssembler. The enrichment of roX2 and H4K16ac (male) on one chromosome depicts full integrity of the assembled X chromosome. **(C)** Overview of HiCAssembler strategy (see methods and Figure S3 for a complete description of the algorithm). The figure displays the iterative progression of the Hi-C assembly strategy as in (Dudchenko et al., 2017) for a small example Hi-C matrix. First, the original scaffolds are split if they contain mis-assemblies and small scaffolds are removed. In each iteration of the Hi-C assembly algorithm scaffolds are joined and oriented to form larger and larger Hi-C scaffolds until chromosome-length assemblies are obtained as shown in the last panel were two separated blocks remain. Afterwards, the small scaffolds that were initially removed are inserted into the Hi-C scaffolds.

**Table 1.**
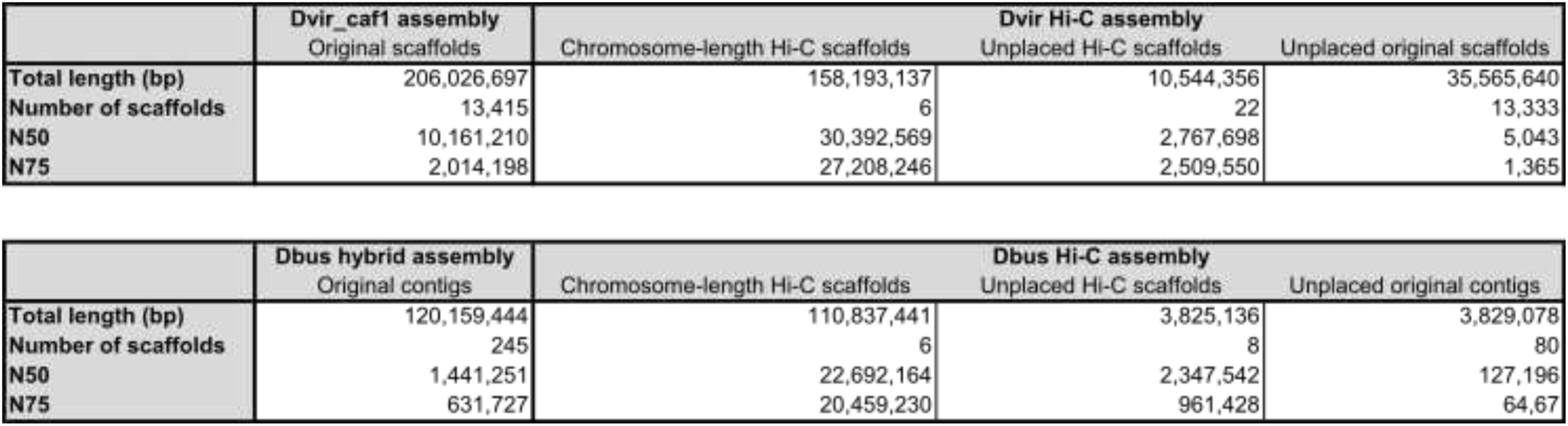
Assembly statistics of the Dvir_caf1 scaffolds and *D. busckii* hybrid contigs (left) that were used as a starting point for the Hi-C scaffolding process into the *D. virilis* and *D. busckii* assemblies (right) reported in this paper. Assembly statistics of the Hi-C assemblies are subdivided into the 6 Muller elements (chromosome-length Hi-C scaffolds), Hi-C scaffolds that were created during the Hi-C assembly process but could not be placed into the 6 Muller elements (Unplaced Hi-C scaffolds) and small original scaffolds/contigs that could not be merged into bigger fragments during the Hi-C scaffolding process (Unplaced original scaffolds/contigs). The reported assembly statistics are: Total length (in bp), total number of scaffolds (referring to contigs or scaffolds before and Hi-C scaffolds after the Hi-C scaffolding process), N50 defined as the length (in bp) at which at least 50 % of the genome sequence is contained in scaffolds of this length and N75 defined as the length (in bp) at which at least 75 % of the genome sequence is contained in scaffolds of this length.

For both species, we used the Hi-C data to identify mis-assemblies as well as to order and orient the pre-assembled contigs/scaffolds into chromosome-length assemblies of 118.5 Gb (*D. busckii*) or 204.3 Gb (*D. virilis*) containing chrX, chr2L, chr2R, chr3L, chr3R and chr4 corresponding to the Muller elements A-F (MULLER and J, 1940) (Figure 1, Table 1 and Figure S2a). Both assemblies contain additional “unplaced Hi-C scaffolds”, which correspond to original contigs/scaffolds that were joined into bigger fragments as well as “unplaced original contigs/scaffolds” that could not be assembled into bigger fragments by Hi-C scaffolding (Table 1).

To produce chromosome-length assemblies of the *D. busckii* and *D. virilis* genomes, our algorithm “HiCAssembler” uses strategies derived from LACHESIS (Korbel and Lee, 2013) and 3D-DNA (Dudchenko et al., 2017), and is freely available at https://github.com/maxplanck-ie/HiCAssembler. In brief, a Hi-C matrix is created by aligning the Hi-C reads to pre-assembled contigs/scaffolds. Then, small fragments (default parameter of 150 kb) are put aside (Figure 1C) and the original contigs/scaffolds are split, if they contain mis-assemblies. In each iteration of the Hi-C assembly algorithm, scaffolds are joined and oriented to form larger Hi-C scaffolds until chromosome-length assemblies are obtained (Figure 1C). Afterwards, initially removed small fragments are inserted into the Hi-C scaffolds (see Figure S3 and the methods section for a detailed description of HiCAssembler, and Table S1 for a comparison with other Hi-C scaffolding tools).

One advantage of the Hi-C assembly process of genomes is that visualization of the Hi-C matrix can be used to detect assembly errors, which appear as discontinuous regions (Figure S3E). The assemblies reported here do not show any conspicuous errors revealed by Hi-C. To further validate this, we generated H4K16ac ChIP-seq of separated male and female larvae and aligned ChIRP-seq of roX2 (Quinn et al., 2016) to our assemblies, as, based on their roles in X chromosome dosage compensation, they are expected to be enriched on the male X. We observe H4K16ac and roX2 enrichment only in one male chromosome-length scaffold, which is present in a single copy in males and two copies in females, and hence, corresponds to the X chromosome (Figure S2C,D). This underscores the validity of our assemblies and the major improvement in terms of continuity over the previous genome assemblies, where roX2 and H4K16ac are scattered across numerous scaffolds (Figure 1A and 1B). We additionally confirmed the quality of our *D. busckii* and *D. virilis* assemblies using Benchmarking Universal Single-Copy Orthologs (BUSCOs) (Waterhouse et al., 2017), which are sets of genes with single-copy orthologs in > 90 % of selected species. BUSCOs can be used to quantitatively measure completeness of genome assemblies (Simão et al., 2015). We tested our genome assemblies with the *Diptera* (odb9) dataset containing 2799 BUSCOs and detected 95.7 % and 98.1 % complete BUSCOs in *D. busckii* and *D. virilis,* respectively (Figure S2E). Such high BUSCO scores emphasize the quality and completeness of our chromosome-length genome assemblies of *D. busckii* and *D. virilis* allowing us to draw valid conclusions about genome topology and evolution in the *Drosophila* genus.

### Conserved TADs are shuffled along the genome during *Drosophila* evolution

Pairwise comparisons between these genome assemblies of *D. virilis* and *D. busckii* with *D. melanogaster* revealed extensive genomic rearrangements between the three species (Figure 2A). We observed that conserved sequences mostly reside on the same chromosomal arms, while only few conserved sequences are found between two different arms. Rearrangements arise without any particular orientation preference. Moreover, shuffling occurs throughout the entire chromosomal arms in the absence of any apparent pattern concerning proximity or distance on the linear DNA sequence.

**Figure 2.**
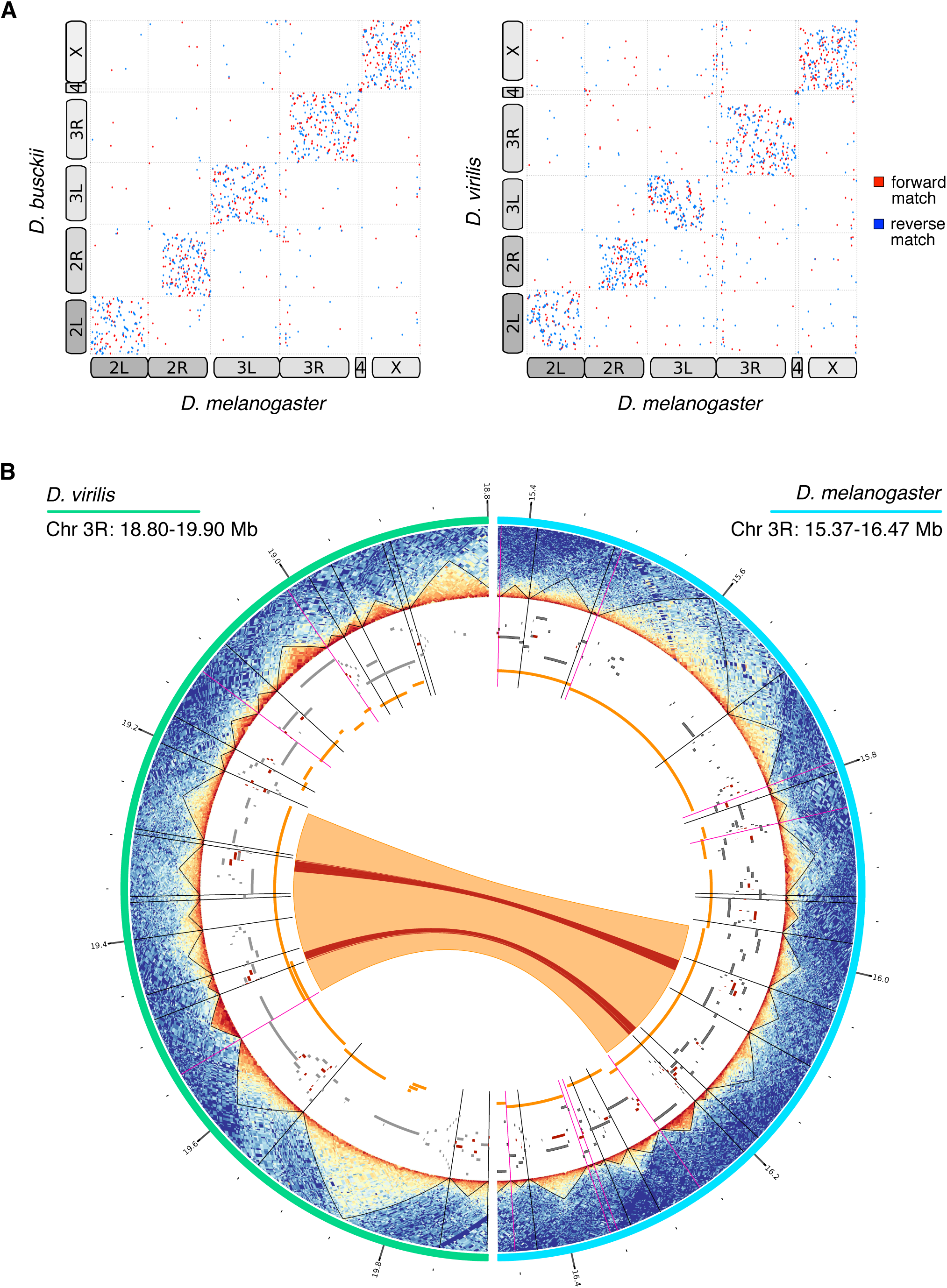
Extensive genome shuffling during *Drosophila* evolution. **(A)** Dotplots showing whole genome alignments between *D. melanogaster* and *D. busckii* or *D. virilis*, respectively. Alignments were performed using Mummer4 (Marçais et al., 2018). Forward matches are shown in red, reverse matches are displayed in blue. Corresponding chromosome arms are indicated with boxes which are displayed connected if chromosome arms are fused in one species. Karyotypes are additionally depicted in Figure S2A. **(B)** Association between TAD boundaries and synteny block breakpoints. From the exterior to the interior of the Circos plot. **Turquoise:** the 18.80 Mb to 19.90 Mb region of the chromosome 3L in *D. virilis.* **Light blue:** the 15.37 Mb to 16.47 Mb of the chromosome 3L in *D. melanogaster*. **Heatmaps**: Hi-C contact heatmaps with TADs displayed as black triangles. **Black radial lines:** TAD boundaries. **Magenta radial lines:** TAD boundaries overlapping with synteny block start or end sites. **Grey blocks:** genes. **Red blocks:** BUSCOs. **Orange blocks:** synteny blocks. **Orange arc:** conserved synteny block between *D. virilis* and *D. melanogaster* in the displayed regions. **Red arcs:** conserved BUSCOs in the displayed regions.

We next analyzed a potential connection between genomic rearrangements occurring during evolution and the 3D architecture of the *D. melanogaster*, *D. busckii* and *D. virilis* genomes. For this, we defined synteny blocks (SBs), which are chains of conserved collinear regions that are used to identify and compare homologous regions between different species. We find, on average, 20 synteny breakpoints per Mb, corresponding to about one breakpoint every six genes. We then compared SBs with two genome topology hierarchies, active/inactive compartments (A/B compartments) and TADs. After obtaining A/B compartments at ∼25 kb resolution from the Hi-C data in all three species (see methods), we correlated the first eigenvector (PC1) of corresponding SBs. We find an r^2^ = 0.45 for *D. melanogaster* and *D. virilis* and r^2^ = 0.42 for *D. melanogaster* and *D. busckii* (Figure S5A). Approximately 75 % of SBs stay within the A or B compartment and 25 % switch between both compartments (Figure S5B). In general, about double the number of SBs lie within the A compartment than B compartment. Therefore, higher order genome topology (A/B compartments), especially the active compartment, appears to be maintained even over 40 million years of genome reshuffling.

We next called TADs in our Hi-C datasets at restriction fragment resolution using HiCExplorer (Ramírez et al., 2018) and validated our TAD calling using several metrics, including a comparison with data that has been sequenced at 10-fold higher sequencing depth (Eagen et al., 2017). We also compared our TAD positions (called using HiCExplorer) with the TAD positions reported by Eagen et al. (called using Arrowhead), which showed great agreement as indicated by ChIP enrichment of the common insulator protein cofactor CP190 (Li et al., 2015). As histone modifications are known to correlate within TADs, we additionally used the H4K16ac ChIP-seq data obtained in all three species to further validate TAD positions (Figure S4). Using these different metrics, we found that our TADs are comparable to the ones reported by Eagen et al. and allow us, despite the lower sequencing depth, to draw valid conclusions about TAD evolution.

When visualizing SBs together with genome topology including TADs, TAD boundaries, BUSCOs and genes (Figure 2B), we noticed that many SB breakpoints overlap with TAD boundaries. Some corresponding SBs also show maintained TAD architecture (see examples in Figure 2B and Figure S5). We therefore quantified the significance of this correlation between genomic rearrangements (i.e. SB breakpoints) and the 3D architecture on a genome-wide level and for this, applied three different methods to validate our findings. First, we computed the overlap of TAD boundaries with SB start and end sites and tested the significance of overlaps, while we used the respective shuffled regions in equivalent analyses as controls (see methods). This analysis revealed, that the overlaps of TAD boundaries and SB breakpoints in all comparisons (Figure 3A) are highly significant (Fisher’s two tailed *p-*value < 2.3e-62), while for shuffled regions, the overlap is decreased to approximately 6 % and not significant (Fisher’s two tailed *p-*value > 0.059, see Table S3). To confirm this result with a second, independent analysis method, we investigated to which extent TADs and SBs overlap in length. To do so, we computed the Jaccard similarity index of TADs and SBs or randomly shuffled SBs as a control. This analysis confirmed a significant difference between the overlap of TADs and SBs in comparison with TADs and randomly shuffled regions (see *p-*values of two-sided Wilcoxon rank-sum test in Figure 3B). As a third approach, we performed pairwise DNA sequence alignments of TADs in the three species and compared their alignment scores to randomly shuffled TADs (Figure 3C). We surmised, that if TADs maintain their integrity during chromosomal rearrangements, their scores obtained from BLASTn (Altschul et al., 1990; Camacho et al., 2009) are expected to be higher compared to random regions. Indeed, we found that the alignment scores (i.e. bitscores) of TADs were significantly higher compared to all controls. This independent sequence-based analysis, which is not taking SBs into account, provided further support for our conclusion that genomic rearrangements in *Drosophila* do not occur randomly but maintain conserved TADs as units.

**Figure 3.**
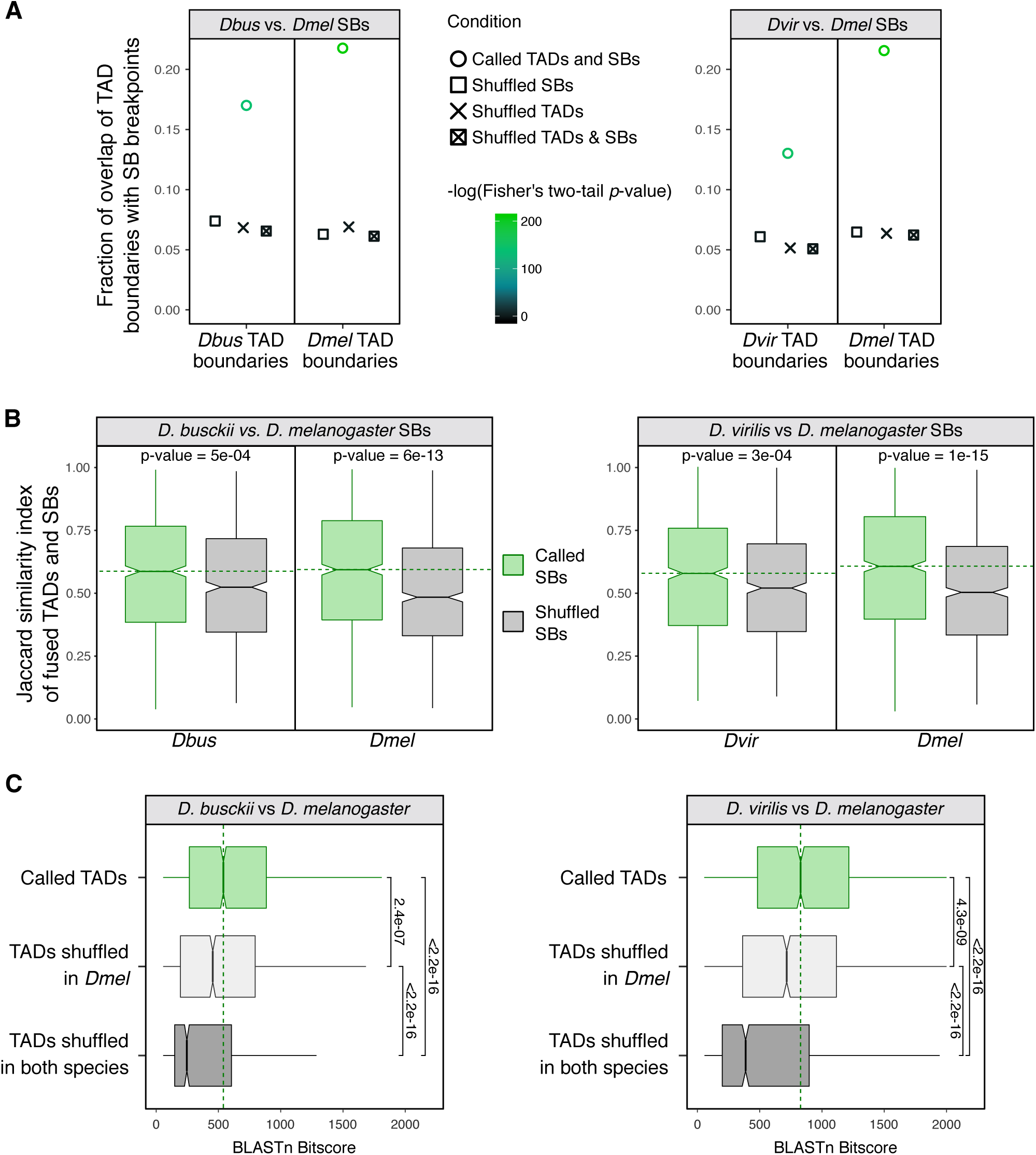
TAD boundaries correlate significantly with synteny block breakpoints. **(A)** Fraction of overlapping extended TAD boundaries with extended synteny block (SB) start and end sites in comparisons of *D. busckii* or *D. virilis* with *D. melanogaster*. The extension is 500 bp in both 5’ and 3’ direction. −log(p-value) of Fisher’s two tailed test is displayed as a color scale of the dots. Overlap and −log(p-value) is shown for boundaries of TADs (n=2209, 2134, 2127 in *D. melanogaster, D. busckii* and *D. virilis*, respectively) and SB breakpoints (n=3726 and 3252 in the *D. melanogaster* vs. *D. virilis* comparison, respectively and 3340 and 2776 in the *D. melanogaster* vs. *D. busckii* comparison, respectively). Overlap with respective shuffled TADs, shuffled SBs and both TADs and SBs shuffled as controls. **(B)** Jaccard similarity index of fused TADs and SBs or respective number of shuffled SBs for *D. busckii* and *D. virilis* compared to *D. melanogaster*. For calculating the Jaccard score, consecutive TADs were fused if a SB overlapped the adjacent TAD by 20 percent or more (see methods). The shuffling of SBs is the same as in (A). The median of called SBs is shown as a green dotted line. *P-*values of two-sided Wilcoxon rank-sum tests are displayed. **(C)** Bitscores of inter-species TAD alignments using BLASTn. TAD to TAD comparisons are displayed in green, TADs shuffled in *D. melanogaster* in light grey, TADs shuffled in both species in dark grey. Shuffling is the same as in (A). The median of called TADs is displayed as a green dotted line and significance was calculated by two-sided Wilcoxon rank-sum test comparisons between the bitscore distributions. All p-values are displayed and significant by using a 0.05 *p*-value threshold.

### Conserved TADs in *Drosophila* are gene-dense and enriched in histone modifications associated with active transcription

Next, we were interested in elucidating whether conserved genome topology, i.e. TADs, relates to particular gene properties, chromatin states or functions. We used a stringent definition to identify conserved TADs among all three species by overlapping TADs with high Jaccard similarity indices and BLASTn bitscores (see methods and Figure 4A). We identified 175 conserved TADs corresponding to approximately 10 % of all TADs, which we compared to an equal number of control TADs (unconserved TADs, see methods) or random genomic regions. Conserved TADs appear significantly bigger compared to all TADs (Figure S6A) and more gene-dense (Figure 4B) compared to unconserved TADs or random regions. The overall length of genes within conserved TADs is similar compared to the other test sets (Figure S6B). 93 % of conserved TADs lie within the active A compartment (Figure 4C), which is a significantly higher proportion compared to unconserved TADs (two-sided two-proportions z-test *p*-value = 0.013) or random regions (*p*-value < 2.2e-16). We next wanted to test, whether conserved TADs are enriched for a particular chromatin state and analyzed them for the five chromatin “colors” reflecting active (yellow and red) and inactive (blue, green and black) states (Filion et al., 2010). Conserved TADs are enriched in the yellow chromatin state associated with broadly expressed genes and the H3K36me3 mark (Figure S6C). As these chromatin “colors” were derived from data obtained in tissue culture cells, we verified this with *in vivo* datasets from fly embryos and larvae. This confirmed that active chromatin marks, such as H3K4me3 and H3K36me3 (embryos, (Celniker et al., 2009)) and H4K16ac (3rd instar larvae, this study) are significantly enriched (bootstrap 95 % CI (confidence interval), *n*=1000) on genes within conserved TADs in comparison with unconserved TADs (Figure 4D,E and Figure S6D). On the other hand, marks as H3K27me3 or the heterochromatin protein 1 alpha (HP1 alpha) associated with gene silencing did not show this trend. Given this association with broadly expressed housekeeping genes (Filion et al., 2010), we checked for the presence of known housekeeping regulators, such as the NSL complex (Lam et al., 2012) and indeed find NSL3 ChIP-seq enrichment at conserved TAD boundaries (Figure 4H, bootstrap 95 % CI (1.23 ± 0.21 in conserved and 0.78 ± 0.23 in unconserved TAD boundaries)). Functional analyses of genes within conserved TADs showed a significant enrichment (Chi-squared test p-value ≤ 0.05) of genes associated with lethal and increased mortality phenotypes upon mutation, which is in line with housekeeping genes encoding for the most fundamental and universal cellular processes (Dickerson et al., 2011) (Figure 4I).

**Figure 4.**
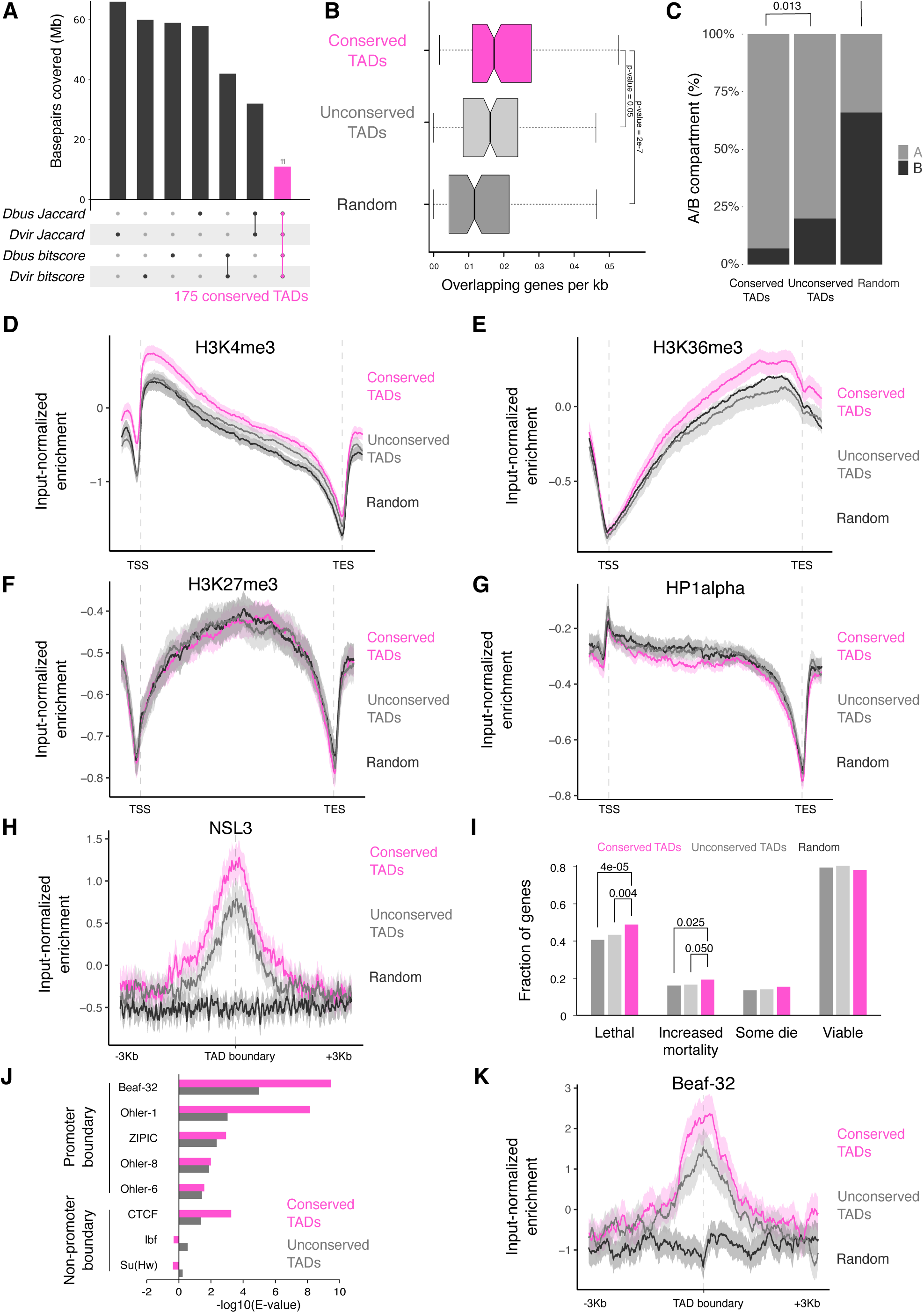
Evolutionary conserved TADs are active gene rich regions comprising essential genes and are demarcated by conserved boundary motifs. **(A)** Definition of conserved TADs between *D. busckii*, *D. virilis* and *D. melanogaster.* TADs with Jaccard similarity index above the median from *D. melanogaster* vs. *D. busckii* and *D. melanogaster* vs. *D. virilis* comparisons were overlapped. Respective overlap of TADs were performed using the bitscores. Afterwards, TADs found in both analyses were compared and the intersect was defined as conserved TADs. Barplots represent the base pair coverage of each subset in the *D. melanogaster* genome. **(B)** Conserved TADs are gene-dense. Genes overlapping with conserved TADs (pink), unconserved TADs (grey) and random genomic regions (dark grey) expressed in number of genes per kb. Equal length of overlapping genes is displayed in Figure S6B as a control. Wilcoxon rank-sum test *p*-values are displayed for comparisons with conserved TADs. **(C)** Percentage of conserved TADs, unconserved TADs and random regions that lie completely in the active (A) or inactive (B) compartment (n = 101, 101, 92). *P*-values were obtained using a two-sided two-proportions z-test. **(D, E, F, G)** Conserved TADs compared to unconserved TADs are significantly enriched in the H3K4me3 **(D)** and H3K36me3 histone marks **(E)** but are not enriched in H3K27me3 **(F)** or HP1 alpha **(G)**. ChIP-seq profiles are from 14-16 h old *D. melanogaster* embryos (Celniker et al., 2009). Log2ratio of H3K4me3, H3K36me3, H3K27me3 and HP1 alpha ChIP-seq reads over input reads along genes (transcription start site (TSS) to transcription end site (TES)) in conserved TADs (pink), unconserved TADs (grey) and random regions (black). ChIP-seq profiles show mean (thick line) and 95 % CI (shadowed area) of input-normalized ChIP-seq enrichment along scaled genes and unscaled 1 kb before the TSS and after the TES. **(H)** Conserved TADs are enriched in the NSL complex member NSL3. Log2ratio of NSL3 (Lam et al., 2012) ChIP-seq reads over input reads at boundaries of conserved TADs (pink), unconserved TADs (grey) and random regions (black) including the 95 % CI (confidence interval) obtained from bootstrapping (n=1000). The NSL3 enrichment at TAD boundaries is significant based on the 95 % CI (1.23 ± 0.21 in conserved and 0.78 ± 0.23 in unconserved TAD boundaries). **(I)** Fraction of genes with lethal, increased mortality, some die or viable phenotypic classes defined in Flybase automatic summaries (genes can be annotated with several phenotypes, see methods). Significant *p*-values (ɑ = 0.05) for genes intersecting conserved TADs are displayed. They were obtained using one-tailed Chi-Squared test to check for proportion differences in two samples. **(J)** Enrichment analysis of promoter and non-promoter boundary motifs at the boundaries of conserved TADs and unconserved TADs in *D. melanogaster.* Beaf-32 shows the highest motif enrichment at conserved TADs in all three species (see Figure S6G). **(K)** Conserved TADs show higher enrichment of Beaf-32 at their boundaries than unconserved TADs by input-normalized ChIP-seq reads (Van Bortle et al., 2014).

We next turned our attention to the boundaries of these conserved TADs and analyzed them for boundary motif enrichments. In *D. melanogaster*, several DNA binding proteins are associated with TAD boundaries, e.g. the Boundary Element Associated Factor-32 (Beaf-32), the motif-1 binding protein (M1BP), the CCCTC-binding factor (CTCF) protein or Suppressor of Hairy-wing (Su(Hw)) (Hug et al., 2017; Ramírez et al., 2018; Sexton et al., 2012; Van Bortle et al., 2014). We previously showed that DNA motifs bound by such factors can be used to predict TAD boundaries in *D. melanogaster* (Ramírez et al., 2018) at high-resolution. Therefore, we analyzed, whether the enrichment of described TAD boundary motifs is conserved in the *Drosophila* species studied here and performed motif enrichment analysis at TAD boundaries in all three species. We focused on motifs previously described in *D. melanogaster* ((Ramírez et al., 2018), see methods) to identify their enrichment at TAD boundaries. We find similar enrichments and comparable E-values for the same boundary motifs in all three species (Figure S6F). We focused more specifically on Beaf-32, as it displayed the lowest E-value at conserved TAD boundaries in all three species (Figure 4J and Figure S6G). Indeed, enrichment of Beaf-32 assessed by ChIP-seq showed higher enrichment at boundaries of conserved TADs in *D. melanogaster* (Figure 4K).

Taken together, our results indicate that irrespective of whether they are conserved or not, TADs in all three species maintain conserved boundary motifs. Despite extensive genomic rearrangements, we find that conserved TADs are gene-dense and enriched in histone modifications associated with active transcription.

### Spatial contacts between high-affinity sites of the dosage compensation complex are conserved during *Drosophila* evolution

The notion that TADs are maintained during evolution points towards an importance of such structures as entities. X chromosome dosage compensation is one example, where adoption of a specialized chromosome architecture has been functionally associated with its chromosome-wide regulation from worms to mammals (Crane et al., 2015; Nora et al., 2012; Ramírez et al., 2015). In flies, this essential process is orchestrated by the MSL complex, which is composed of the noncoding RNAs roX1 and/or roX2, as well as the proteins MSL1, MSL2, MSL3, MLE and MOF (Kuroda et al., 2016; Samata and Akhtar, 2018). In agreement with earlier findings (Meisel et al., 2012; Quinn et al., 2016), immunostainings of male and female polytene chromosomes showed a strong male-specific enrichment of MOF on the X chromosome of *D. melanogaster, D. busckii* and *D. virilis* (Figure 5A and Figure S7A). MSL recruitment to the X chromosome occurs at special binding sites termed high-affinity sites (HAS), which are particularly enriched for MSL2, MLE and roX1/2 and cluster together in space (Alekseyenko et al., 2008; Ramírez et al., 2015; Schauer et al., 2017; Straub et al., 2013; Valsecchi et al., 2018). The presence of HAS sequences and their enrichment of roX appears to be a conserved feature of the X within the *Drosophila* genus (Alekseyenko et al., 2013; Ellison and Bachtrog, 2013; Quinn et al., 2016). Given our finding of extensive genome rearrangements, we were interested how shuffling of the X chromosome impacts dosage compensation and in particular the clustering of HAS into a “dosage compensation hub” in 3D. We defined a comparable set of high-confidence HAS in all three species using roX2 ChIRP-seq data (Quinn et al., 2016) and then analyzed our Hi-C data for the conservation of genome-topology at those sites (see methods). By using overlapping BUSCOs, we identified corresponding HAS in the three species, which revealed that they have substantially changed their relative position within the X chromosome (Figure 5B,C). Our chromosome-length assemblies also allowed us to specifically inspect genes that switched between autosomes and the X chromosome in between species. For example, *MED20* moved from chromosome 2L in *D. melanogaster* to the X chromosome in *D. virilis*, where it now resides within a H4K16ac-positive domain (Figure 5D). Another example of a gene that moved in the opposite direction, namely from the X chromosome in *D. melanogaster* to chromosome 2L in *D. virilis,* is *mei-41* (Figure S7B). This gene resides in a H4K16ac-positive domain on the X chromosome and only retains a H4K16ac promoter-peak when moved to the autosome. Another example of a gene that moved from chromosome 2L in *D. melanogaster* to the X chromosome in *D. virilis* and there gained a species-specific HAS is the gene *dim gamma-tubulin 3* (Figure S7C).

**Figure 5.**
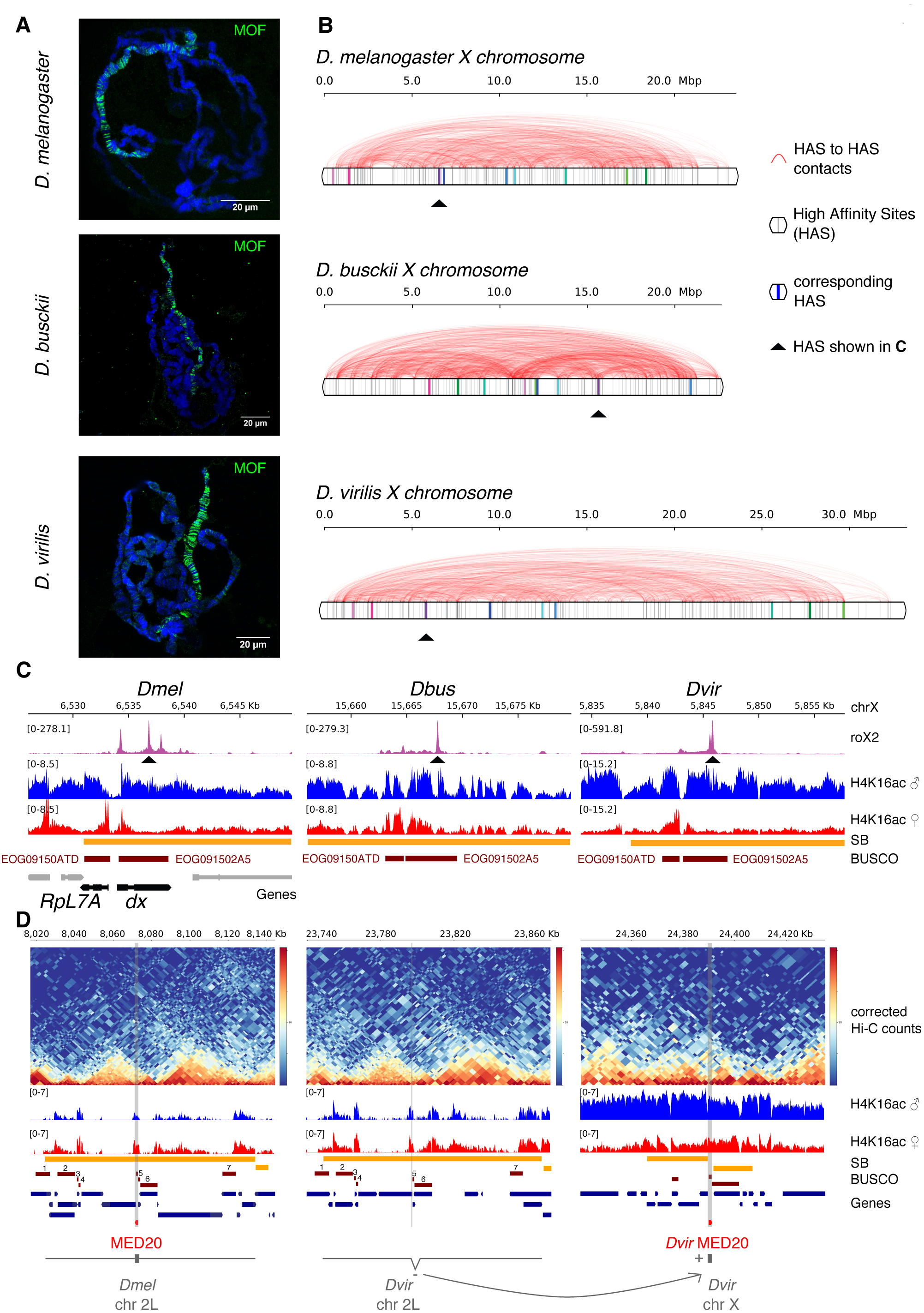
Binding sites of the dosage compensation complex are shuffled along the X chromosome in *D. melanogaster, D. busckii* and *D. virilis*. **(A)** Immunostaining of male polytene chromosomes with MOF antibody (green) in *D. melanogaster, D. busckii* and *D. virilis*. DNA is counterstained with Hoechst (blue). Scale bars, 20µm. Immunostaining of female polytene chromosomes are shown in Figure S7. **(B)** Position (gray vertical bars) and obs/exp Hi-C contacts (red arcs) between high-confidence roX2 sites (HAS) along the entire X chromosome in *D. melanogaster, D. busckii* and *D. virilis*. **(C)** Example of one corresponding HAS as indicated by the red arrows in (B). Coverage of roX2 ChIRP-seq reads, H4K16ac ChIP-seq reads from separated female and male 3rd instar larvae, SBs, BUSCOs and genes annotated in *D. melanogaster*, highlighting *RpL7A* and *dx*, two genes with phenotypic classes related to viability reduction corresponding respectively to the BUSCOs EOG09150ATD and EOG091502A5 in all three species. **(D)** Example gene that moved between an autosome and the X chromosome when comparing *D. melanogaster* and *D. virilis*. MED20 is localized on chromosome 2L in *D. melanogaster* but on chromosome X in *D. virilis* (*Dvir* GJ18844). The surrounding genes on this SB on chromosome 2L maintained the same order (see corresponding BUSCOs numbered from 1-7 and gene track). MED20 in *D. virilis* (*Dvir* GJ18844) is localized on chromosome X in between two surrounding SBs, within a H4K16ac domain (male). Two additional examples are shown in Figure S7B,C.

In light of these genome-wide genomic rearrangements, we asked whether spatial contacts between HAS could possibly be conserved. Indeed, aggregate plots revealed an enrichment of Hi-C contacts for HAS but not random control regions (Figure 6A) in all three species, highlighting the maintenance of these interactions despite extensive shuffling of the genomes. As HAS are enriched near TAD boundaries in *D. melanogaster* (Ramírez et al., 2015), we wanted to analyze whether this also holds true in other *Drosophila* species and calculated the distance of the 250 high-confidence HAS to the closest TAD boundary (Figure 6B). Indeed, *D. virilis* and *D. busckii* HAS, but not random regions, are also enriched in the vicinity of TAD boundaries. In agreement with earlier observations in *D. melanogaster* (Hug et al., 2017), we were also able to identify enriched TAD boundary contacts in *D. busckii* and *D. virilis* (Figure 6C). Because HAS tend to be in the proximity of boundaries, we therefore wanted to test if HAS-HAS contacts were simply driven by their “closeness” to boundaries. For this, we defined a set of alternative, non-HAS genomic positions around TAD boundaries (defined as “mirrored HAS”), which have the same distance than HAS to TAD boundaries but are located on the opposite site of the closest TAD boundary. Interestingly, we do not find enriched contacts at these “mirrored HAS” (Figure 6C), while gene expression levels between HAS-genes (n=209) and mirrored HAS-genes (n=181) were similar (Figure 6D). Furthermore, non-HAS genes, which are expressed equally strong or higher than HAS genes, showed lower Hi-C contacts in comparison with HAS (Figure 6E and Figure S7D,E). This suggests that enriched contacts between HAS are not a mere consequence of closeness to TAD boundaries or the expression level of their associated genes.

**Figure 6.**
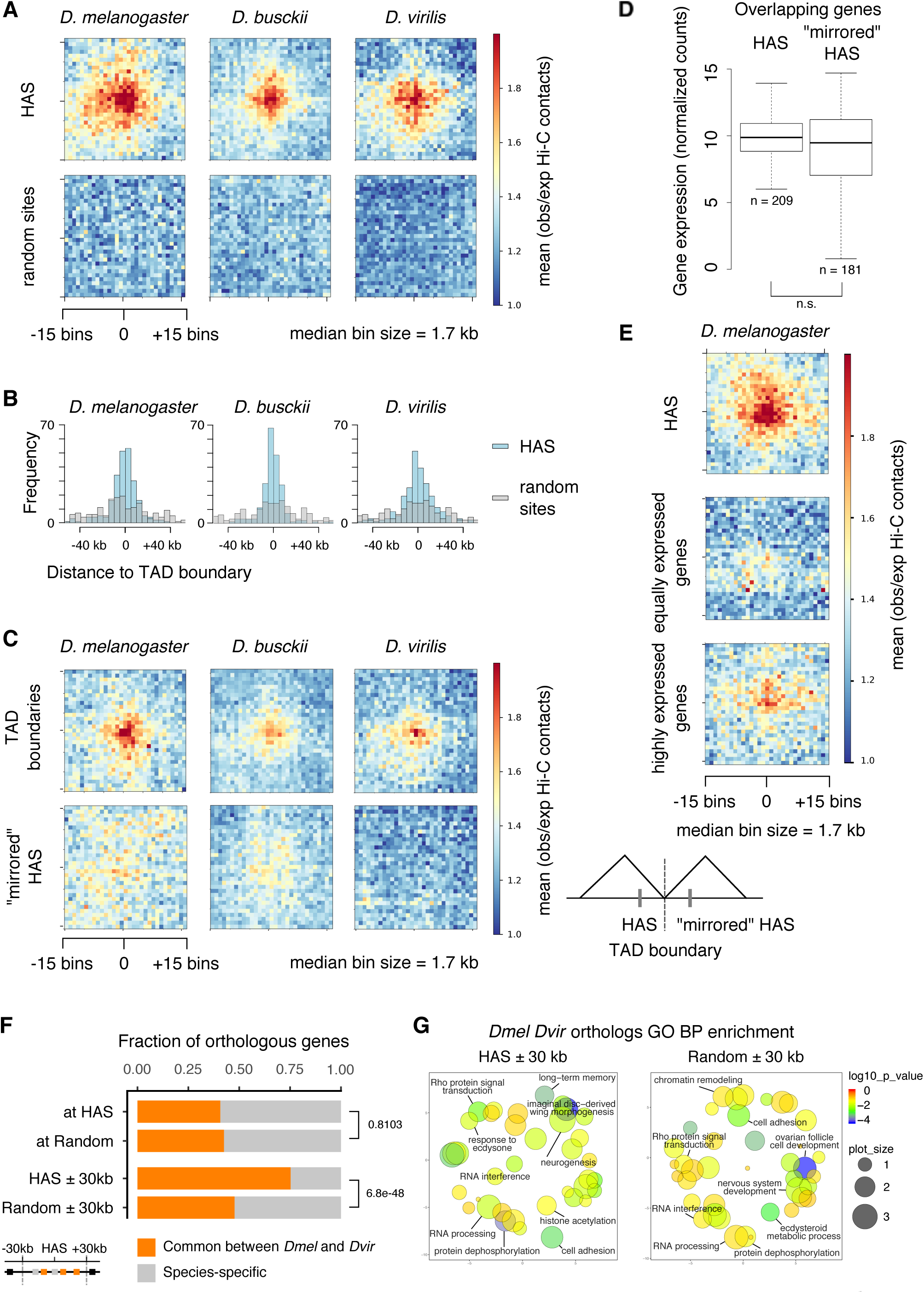
Enriched Hi-C contacts between binding sites of the dosage compensation complex are maintained throughout *Drosophila* evolution despite genome shuffling. **(A)** Aggregate Hi-C matrices around pairwise HAS-HAS contacts in *D. melanogaster, D. busckii* and *D. virilis* compared to random pairwise contacts. Displayed are the mean observed over expected contacts ratios of corrected Hi-C matrices with a ∼1.7 kb bin size of approximately 250 HAS that are on the X chromosome (n=246, 213 and 247 in *D. melanogaster, D. busckii* and *D. virilis*, respectively) or a respective number of random regions on the X chromosome. **(B)** Distance of HAS used in (A) to closest TAD boundary compared to the respective number of random sites. **(C)** Aggregate Hi-C matrices centered on TAD boundaries with the lowest insulation score on the X chromosome in *D. melanogaster, D. busckii* and *D. virilis* (n=246, 213 and 247 in *D. melanogaster, D. busckii* and *D. virilis*, respectively), and HAS mirrored at their closest TAD boundary (“mirrored” HAS). TAD boundaries show enriched contacts but “mirrored” HAS show no enriched contacts. **(D)** Gene expression (normalized counts) of genes overlapping with HAS compared to “mirrored” HAS is not significantly different (n.s.) by Wilcoxon rank-sum test. Comparison of gene expression was performed using library size normalized RNA-seq counts from 14-20 h aged embryos from modENCODE datasets obtained from (Ramírez et al., 2018) and also available on the chorogenome web server (http://chorogenome.ie-freiburg.mpg.de/). **(E)** Aggregate Hi-C matrices around HAS-HAS or TSS-TSS contacts of genes with equally high expression as genes overlapping with HAS or highly expressed genes in *D. melanogaster* (n = 209). Underlying gene expression values are shown in Figure S7E. **(F)** Fraction of total orthologous genes (grey) and orthologous genes in common between *D. melanogaster* and *D. busckii* (orange) at HAS (n=195), at random positions defined in the X chromosome A (active) compartment (n=175), at HAS extended by 30 kb in 5’ and 3’ direction (n=1380), and at 60 kb random regions defined in the X chromosome A (active) compartment (n=838). Reported values are calculated in *D. melanogaster*, n is equal to the total number of orthologs between the two species for each category. Two-sided two-proportions z-test *p*-values are shown on the right of the bar plot. Orthologs between *D. melanogaster* and *D. virilis* were retrieved from flybase. **(G)** Gene Ontology of Biological Processes (GO BP) enrichment of genes overlapping HAS and random active regions extended by 30 kb in 5’ and 3’ direction. Only genes with orthologs between *D. melanogaster* and *D. virilis* (*Dmel Dvir* orthologs) were considered.

We next wanted to compare orthologous genes at HAS between *D. melanogaster* and *D. virilis.* Using HAS positions in the *D. melanogaster* genome, comparison of orthologs showed that about 40 % of genes at HAS are in common between the two species (Figure 6F). The fraction of common genes is similar to what would be expected by chance in the A compartment (*p*-value = 0.81) where we detected about 42 % of common orthologs between *D. melanogaster* and *D. virilis*. We compared HAS to genes in the A compartment because most HAS fall within the A compartment. Yet, when comparing genes in the vicinity of HAS (± 30 kb), we find about 73 % conserved orthologous genes, which is significantly different from active random regions showing 46 % conserved orthologous genes (*p*-value < 2.2e-16). We choose 30 kb extensions around HAS because distances between HAS are about 60 kb in all three species, and because gene expression was shown to be affected up to this distance upon mutation of roX1 and roX2 (Kim et al., 2018). The respective analysis in *D. virilis* showed similar trends (*p*-value = 2.3e-4 at HAS and 6.2e-21 ± 30 kb around HAS). Since HAS themselves are not enriched at genes that are evolutionary conserved while the genes in their vicinity (30 kb) are, we asked whether individual conserved genes in the vicinity of HAS rely on the same HAS for compensation or not. To test this, we determined each HAS identity between *D. melanogaster* and *D. virilis* by finding pairs of HAS with the highest fraction of common orthologous genes between both species compared to all orthologous genes in their vicinity (± 30 kb) (Figure S7F). We repeated this analysis for 100 sets of random 60 kb regions in *D. melanogaster* and *D. virilis* and report no significant difference to extended HAS (Wilcoxon test *p*-value = 0.99 after adjustment for multiple comparison by Benjamini-Hochberg procedure). This suggests that compensated genes stay in the vicinity of any HAS but not necessarily the same HAS over evolution. GO term analysis of genes around HAS showed similar enriched functions compared to respective random regions (Figure 6G). This suggests that compensated genes are evolutionary conserved and involved in diverse and essential biological functions.

Taken together, our data strongly suggests that regulatory relevant genome 3D architecture, for example for X chromosome regulation, is maintained during evolution independently of chromosome rearrangements.

## Discussion

Here, we analyzed the impact of genomic rearrangements on genome topology using the *Drosophila* genus and X chromosome dosage compensation as a model system. After generating Hi-C assisted chromosome-length assemblies of the *D. busckii* and *D. virilis* genomes, we identified and characterized conserved TADs as evolutionary maintained units in three *Drosophila* species and further analyzed the conserved spatial network of HAS contacts on the X chromosome.

Hi-C scaffolding has been shown to produce high-quality assemblies with continuity up to chromosome-length in various species such as the mosquito *Aedes aegypti*, the domestic goat or barley and in some cases by integrating several data types such as optical mapping or 3rd generation sequencing data (Bickhart et al., 2017; Burton et al., 2013; Dudchenko et al., 2017; Korbel and Lee, 2013; Marie-Nelly et al., 2014; Mascher et al., 2017). In our study, we combined *de novo* assembled contigs of homogeneous size - in the case of *D. busckii* - or using published scaffolds of diverse sizes as a starting point - in the case of *D. virilis* - with Hi-C data to generate chromosome-length genome assemblies. We developed HiCAssembler as a free, open source and user-friendly Hi-C scaffolding tool that automatically visualizes the scaffolding process, which thus allows easy fine-tuning of parameters and correction of assembly errors. The Hi-C libraries for scaffolding were sequenced at a coverage as little as 17X for *D. busckii* and 19X for *D. virilis.* Hence, our approach allows to generate high quality chromosome-length assemblies in an efficient and cost-effective manner. Contrary to our approach, genome assemblies only produced by long-read technologies such as PacBio require high coverage to produce high quality and contiguous assemblies, which remains cost-inefficient. Chakraborty et al. for example used a 121X PacBio read coverage to reach a N50 of approximately 20 Mb corresponding to chromosome length in *D. melanogaster* and tested by downsampling that lower coverage assemblies only yield fragmented assemblies (Chakraborty et al., 2016, 2018).

Despite the advantages of Hi-C scaffolding we note that both generated assemblies in our study still contain a small fraction of unplaced fragments (also see “Unplaced Hi-C scaffolds” and “Unplaced original contigs/scaffolds” (Table 1)). These fragments are repetitive and/or too small to be placed with enough confidence in the Muller elements without introducing assembly errors using Hi-C scaffolding. Long read sequencing can aid in the assembly of such repetitive regions, and accordingly, we note that the fraction of unplaced scaffolds in our *D. busckii* genome assembly is lower than in the *D. virilis* assembly. Yet, all Hi-C based genome scaffolding methods including ours only consider mappable regions of the genome and hence, our assemblies do not contain highly repetitive and unmappable regions of the genome. While the analyses discussed hereafter do not integrate repetitive regions and are thus not affected by the omission of such regions, it will be important to further improve genome assemblies for studying genome evolution of repeats. Nevertheless, our two chromosome-length genome assemblies of *D. virilis* and *D. busckii* show an unprecedented continuity compared to the previous assemblies of these species’ genomes and thus constitute a valuable resource for further genetic and evolutionary studies in *Drosophila*.

Our analyses reveal that genomic rearrangements during evolution occurred mostly within the entire length of the same chromosome arms (Figure 2A). This observation is in accordance with previous reports, stating that orthologous genes are typically found on the same Muller element (Drosophila 12 Genomes Consortium et al., 2007). Similarly, rearrangements preferentially occur within the same chromosome arms in the mosquito species *Aedes aegypti, Culex quinquefasciatus* and *Anopheles gambiae,* which are separated by about 150 to 200 million years of evolution (Dudchenko et al., 2017).

We took advantage of our chromosome-length assemblies and the associated chromosome conformation data in order to comprehensively define and characterize conserved sequences between the *D. melanogaster*, *D. busckii* and *D. virilis* genomes. We identified synteny blocks (SBs) between the three *Drosophila* species and found conservation of active and inactive (A/B) compartments in these regions. Moreover, we report significant overlap of synteny breakpoints with TAD boundaries. This observation suggests that the positions of TAD boundaries are associated with the position of synteny breakpoints and a fraction of TADs appear maintained as evolutionary conserved units in *Drosophila*. Moreover, we demonstrated that conserved TADs are mainly found in the active compartment, showing that genome shuffling and TAD structure rearrangements are more selected against in the active compartment than in the inactive compartment during evolution. Genes within conserved TADs are involved in essential functions and are enriched for active histone modifications. This functional relevance and/or their regulatory surroundings may explain why they have been maintained during evolution. We hypothesize that ancient TADs are conserved because rearrangements are negatively favoured during evolution within these domains containing active, gene-dense regions important for maintaining cell integrity.

Our genome-wide analyses suggest that conserved TADs may indeed explain a fraction of the non-random distribution of synteny breakpoints. However, other factors than TADs are involved in the determination of these breakpoints, as a significant fraction cannot be associated with a conserved TAD structure, at least with our analyses. In mammals, retrotransposition was shown to create new CTCF-binding sites by repeat element expansions (Schmidt et al., 2012). In principle, such a mechanism could be also operating in flies, where transposable elements might shape genome topology, TAD boundaries and breakpoints. We note that conserved TADs are demarcated by stronger but not different sequence motifs *per se* in comparison with unconserved TADs. This could indicate that TAD boundaries with strong motifs are more open and accessible and thus break or rearrange easier than sites within the TADs around them. Further studies are required to elucidate, if there is any preference of chromosome breakage at certain sequences, e.g. during germ cell formation, and how such rearranged chromosomes are inherited and selected against later on during development, in particular if regulatory topology and thus possibly gene expression is affected.

The occurrence of evolutionary rearrangements at TAD boundaries was also described in a gibbon-human comparison (Lazar et al., 2018). This study identified 67 rearrangements between both species within an evolutionary distance of approximately 19 million years. The *Drosophila* species analyzed in our study cover approximately 40 million years of evolution and multiple subgenera (Robe et al., 2010; Russo et al., 1995). We observe that Drosophila genomes are apparently extremely rearranged with, on average, 20 synteny breakpoints per Mb despite being highly gene-dense. This high number of synteny breakpoints is in agreement with earlier reports (Bhutkar et al., 2008) and may be ascribed to the short generation time of *Drosophila* (Thomas et al., 2010) compared with mammalian species. Therefore, the comparison of *Drosophila* species is extremely powerful in elucidating fundamental properties of conserved TADs despite thousands of genomic rearrangements and allows characterization of their features.

Genome topology is involved in facilitating the spreading of the dosage compensation complex on the X chromosome by spatial proximity of HAS (Ramírez et al., 2015; Schauer et al., 2017). This 3D conformation is maintained during evolution in *Drosophila*, as we observe enriched HAS contacts in all three analyzed species. Taking into consideration that genomes are completely shuffled during evolution, the conservation of dosage compensation (Alekseyenko et al., 2013; Quinn et al., 2016) and the recognition of the X chromosome specifically is indeed remarkable. This strongly suggests that enriched contacts between HAS are conserved in the *Drosophila* genus because of their functional importance for dosage compensation. Indeed, our analyses imply that HAS do not exhibit spatial proximity simply because of their closeness to TAD boundaries or gene activity, but rather intrinsically provide this property. Yet, this seems not to involve male-specific factors, as such enriched contacts are found in both males and females (Ramírez et al., 2015; Schauer et al., 2017).

Several models exist that describe spreading of the dosage compensation complex. Conserved HAS were shown to be more evenly spaced on the linear X chromosome than expected by chance which could allow easier and more effective spreading of the dosage compensation complex (Quinn et al., 2016). Integrating our findings about the X chromosomal 3D topology, enriched contacts between HAS are likely to reflect their connection into a spatial network that includes the entire X chromosomal territory, in which the dosage compensation complex is confined. This idea of spatially co-localized binding sites was suggested for transcription factors and enhancers using imaging-based techniques (Liu et al., 2014; Pernuš and Langowski, 2015), biophysical modelling (Brackley et al., 2012; Malin et al., 2015) or Hi-C and single cell Hi-C data (Ma et al., 2018). In summary, these studies demonstrate that spatial proximity of binding sites can facilitate diffusion and increase local concentration of factors that need to be distributed along the genome. The connections in such networks can be variable between single cells (Finn et al., 2019; Ma et al., 2018). Thus, the spatial network of HAS could theoretically enhance spreading of the dosage compensation complex along the entire X. Further studies are required to understand the molecular mechanism of 3D interaction site formation of HAS. Another hint towards the importance of spreading but not necessary the location of individual HAS themselves is that genes in the vicinity of HAS are more conserved than genes overlapping with HAS. Genes that require dosage compensation could be located around HAS where the dosage compensation complex binds and subsequently spreads into its surroundings. This suggests that the spatial proximity of HAS and the spreading of the dosage compensation complex from them is more important and conserved with respect to the dosage compensation function than the absolute positioning and order of individual HAS. Thus, HAS seem interchangeable and contributing equivalently to dosage compensation across evolution. This could be similar for other binding sites, important for essential regulatory mechanisms, that display spatial proximity such as enhancers, although further studies are required to test this hypothesis.

In summary, the Hi-C guided *Drosophila* assemblies and the strategies for comparing chromatin conformation data between species presented in our study, provide insights into genome topology evolution. In particular, our finding of evolutionary stability of entire regulatory units on chromosomes and topology including a full chromosome, despite genome-shuffling, may be an important step towards a further understanding, in how changes and mutations affecting gene topologies may impact on essential cellular processes in all eukaryotic kingdoms.

## Acknowledgements

We thank Aline Gaub for help with polytene chromosome squashes, Vivek Bhardwaj for helpful discussions and Plamen Georgiev for critically reading the manuscript. We also thank the Deep Sequencing Facility (Ulrike Boenisch and team) and the Bioinformatics Facility (Thomas Manke and team) of MPI-IE for their help. G.Ri was funded by French National Institute for Agricultural Research Young Scientist Contract (INRA CJS). CIKV was supported by a Human Frontier Science Program (HFSP) long-term fellowship (000233/2014-L). L.A. was funded by the German Epigenome Programme “DEEP” (BMBF project number 01KU1216G).This work was supported by DFG funded CRC992, CRC1140 awarded to A.A.

## Author Contributions

G.Re., F.R., S.T. and A.A. conceptualized the study. S.T. arranged PacBio sequencing and initiated Hi-C experiments in fly embryos. G.Re. performed fly embryo sample collection. G.Re. performed Hi-C experiments with the help of L.A.. G.Re. and F.R. assembled the *D. busckii* and *D. virilis* genomes. F.R. developed HiCAssembler. G.Re., G.Ri. and F.R. analyzed all datasets, conceptualized and performed all data analyses and their graphical representations, and interpreted the biological significance of the results. C.I.K.V. performed H4K16ac ChIP-seq experiments and contributed to the analysis and interpretation of the data. G.Re. performed polytene chromosome squashes and microscopy. F.R. and A.A. mentored and supervised the research. The manuscript was fully drafted by G.Re., written and edited by G.Re., C.I.K.V., G.Ri., F.R., and A.A. All authors reviewed, edited, and approved the paper.

## Code availability

HiCAssembler is freely available at https://github.com/maxplanck-ie/HiCAssembler.

## Data availability

The raw PacBio and Hi-C data have been deposited to the NCBI Sequence Read Archive (SRA) and can be found under accession SRR7029387-SRR7029398. ChIP-seq and processed Hi-C data including both genome assemblies have been deposited to Gene Expression Omnibus (GEO) and can be found under accession GSE120752.

## Methods

### *D. melanogaster*, *D. virilis* and *D. busckii* fly lines

The following fly lines were used for experiments: *D. melanogaster* (*w^1118^* Oregon R *Drosophila melanogaste*r), *D. virilis* (*Drosophila virilis*, San Diego stock center, stock number: 15010-1051.00) and *D. busckii* (*Drosophila busckii*, San Diego stock center, stock number 13000-0081.31).

*D. melanogaster* and *D. virilis* flies were maintained at room temperature on standard cornmeal-molasses medium. *D. busckii* flies were additionally fed with instant *Drosophila* medium (Formula 4-24, Carolina Biological Supply Company, catalog number 173202) mixed with instant potato powder on top of the standard *Drosophila* medium.

### Fly embryo collection and fixation for *in situ* Hi-C

Flies were transferred into collection cages at 25 °C at least one day before embryo collection. Pre-lays were done for 2 h and fly embryos were collected on apple juice plates with yeast for 2 h and then aged at 25 °C until they reached developmental stage 15-16. Because embryogenesis timing differs across species (Kuntz and Eisen, 2014) we collected 16-18 h old *D. melanogaster*, 21-23 h old *D. virilis* and 19-21 h old *D. busckii* embryos (see Figure S1). Hi-C data from 21-23 h old *D. busckii* embryos was used for the genome assembly of *D. busckii*. Because no differences in the 3D chromatin conformation were found between males and females, we used mixed embryos in our experiments (Ramírez et al., 2015; Schauer et al., 2017).

*D. melanogaster* and *D.virilis* embryos were dechorionated, washed and fixed for 15 min in 5 mL of 1 % methanol-free formaldehyde in PBS with 5 mL heptane while shaking. Fixation was stopped by adding glycine up to a final concentration of 0.25 M and incubating for 5 min. The fixation solution was removed and embryos were washed two times in 0.1 % Triton-X in PBS for 10 min. Supernatant was removed and samples stored at −80°C. To maintain the integrity of all nuclei, including those far from the embryo surface, we fixed *D. melanogaster* and *D. virilis* embryos again while breaking them into smaller fragments in a 1 mL dounce homogenizer using 1 % methanol-free formaldehyde in serum-free Schneider’s medium supplemented with 0.5 % NP-40 for 10 min at room temperature. Fixation was quenched by adding glycine up to a final concentration of 0.125 M and immediate pelleting of fly embryo fragments at 1000 g for 5 min. Samples were washed in PBS and then kept on ice for nuclei extraction (see next paragraph). *D. busckii* embryos are smaller than embryos from the other two fly species (Gregor et al., 2005). Fixation using the above described procedure lead to loss of many embryos during fixation. Therefore, *D. busckii* fly embryos were dechorionated and directly fixed while breaking them into smaller fragments in a 1 mL dounce homogenizer as described above.

### *In situ* Hi-C of *D. melanogaster*, *D. virilis* and *D. busckii* embryos

*In situ* Hi-C experiments were performed using a modified version of the *in situ* Hi-C protocol (Rao et al., 2014) described in (Ramírez et al., 2018). Nuclei were extracted by resuspension in 1 mL ice-cold lysis buffer (10 mM Tris-HCl, pH 8, 10 mM NaCl, 0.2 % IGEPAL CA-630) and sonication following the standardized nuclear extraction NEXON protocol (Arrigoni et al., 2015) (Covaris E220 sonicator, settings: 75 W peak power, 2 % duty factor, 200 cycles/burst, for 15-30 sec until about 70 % of intact nuclei were released). Samples were filtered through a 30 µm filter to remove bigger embryo fragments. From this step on we followed the protocol described in (Ramírez et al., 2018). Nuclei were digested using DpnII (NEB, R0543M, 150 U/ sample). To increase the fraction of valid Hi-C reads (see Figure S1), dangling ends were removed after purification of Hi-C ligated DNA in samples from *D. busckii* using 5U of T4 DNA polymerase for 30 min at 20°C with addition of 25µM dGTP. After biotin pull-down, 50 ng of DNA bound to beads was used for library preparation and libraries were sequenced paired-end, with a read length of 75 bp, on a Illumina HiSeq 3000 or Illumina NextSeq500 machine. Table S2 provides the numbers of sequenced and filtered valid reads of all Hi-C samples.

### *De novo D. busckii* hybrid contig assembly

To assemble the *D. busckii* genome we followed a hybrid approach (Figure 1A) in which we combined short Illumina and long PacBio reads. Illumina reads of female flies were obtained from (Zhou and Bachtrog, 2015) and (Vicoso and Bachtrog, 2015) accession codes SRR1795010, SRR1794619, SRR1794616, SRR1794617, SRR1794614 and SRR826809. These paired-end reads of whole genome sequencing data were trimmed for adapters and sequencing quality using Trim Galore (v0.4.0). All reads (68 bp average length, 18.8 Gb, 156X coverage) were merged like single-end data and assembled into contigs using SparseAssembler v20160205 (Ye et al., 2012) with parameters ‘*k 51 g 15 NodeCovTh 1 EdgeCovTh 0 TrimN 2 GS 240000000*’. We did not include the paired-end information from Illumina reads using SparseAssembler to reduce errors introduced by heuristics usually applied in short read assembly as gap closing or scaffolding. Short read assembly resulted in 32,010 contigs with N50 of 8.9 kb.

PacBio reads of genomic DNA were generated by GATC Biotech from 100 adult female *D. busckii* flies. PacBio reads were sequenced on a RS II system using P6 chemistry. In total, 2.7 Gb PacBio data (20X coverage) were obtained with a mean polymerase read length of 8.2 kb and a mean subread length of 5.7 kb. The FM-index Long Read Corrector FMLRC (Wang et al., 2018) was used to reduce sequencing errors of the PacBio reads by using short Illumina reads. In a second assembly step, PacBio reads were integrated by aligning and overlapping the previously generated high-confidence contigs to the much longer but error-prone PacBio reads using DBG2OLC v20160205 (Ye et al., 2016) with parameters ‘*LD 0 k 17 AdaptiveTh 0.01 KmerCov 2 MinOverlap 20 RemoveChimera 1 ChimeraTh 2*’. DBG2OLC assembly resulted in a total assembly length of 120,159,444 bp consisting of 245 contigs with N50 of 1,441,251 bp (Table 1). This hybrid assembly approach takes advantage of highly accurate Illumina reads and long 3rd generation sequencing reads. The next paragraph describes chromosome-length scaffolding of these contigs using Hi-C data.

We have tested several additional assembly methods for contig assembly that allow combining PacBio and Illumina reads or PacBio data alone, namely Canu (Koren et al., 2016), Miniasm (Li, 2016), Spades (Bankevich et al., 2012) and Masurca (Zimin et al., 2013). After Hi-C scaffolding, we evaluated the total assembly length, mapping rate of Hi-C data to the assemblies as well as frequency of obvious mis-assemblies by visual inspection of the automatically generate Hi-C matrices from HiCAssembler. We concluded that the assembly provided by SparseAssembler in combination with DBG2OLC was the best and generated subsequent assemblies by fine tuning the parameters to reduce mis-assemblies that are clearly visible as part of the assembly of the contigs using Hi-C.

### Hi-C assembly algorithm

To assemble the *D. busckii* and *D. virilis* genomes we used an iterative scaffolding strategy similar to 3D-DNA (Dudchenko et al., 2017). To determine the order of scaffolds we used a maximum spanning tree as in LACHESIS (Korbel and Lee, 2013). Our algorithm is open source, easy to install and to use and is freely available at https://github.com/maxplanck-ie/HiCAssembler.

Our assembly of chromosome-length Hi-C scaffolds from pre-assembled short contigs/scaffolds consists of the following steps (in the text we will use ‘scaffolds’ to refer to pre-assembled short contigs or already available scaffolds):

#### Creation of corrected Hi-C contact matrix

##### Read mapping

Reads are aligned to the pre-assembled scaffolds, each mate is aligned separately using BWA MEM (Li, 2013) with parameters -A1 -B4 -E50 -L0 (which promote a read to be split instead of adding a gap).

##### Creation of Hi-C contact matrix

’*hicBuildMatrix*’ from HiCExplorer (Ramírez et al., 2018) is used to to compute the Hi-C contact matrix after filtering out low-quality reads. They consist of reads that are mapping to several repetitive regions, that are not close to restriction sites, that did not re-ligate (dangling ends), self-circles and same-fragment reads. Information about read filtering during matrix creation is provided in a quality control (QC) summary of each Hi-C sample. Bin size is set to restriction fragment length.

##### Matrix correction

The total number of reads that are assigned to each bin is calculated. Bins having zero or low number of reads as well as bins having read counts over 1.6 median absolute deviation scores (MAD-score) are removed. The elimination of bins with a low number of reads avoids amplification of signal from these bins during the correction using matrix balancing. The elimination of bins with a MAD *z*-score of 1.6 or larger aims to reduce bins containing collapsed repetitive regions due to assembly errors. Collapsed repetitive regions refer to genomic repetitions that appear as unique in pre-assembled scaffolds which can be identified in some cases by a high coverage. After bin filtering, the matrix is corrected using iterative correction (Imakaev et al., 2012) implemented in the ‘*hicCorrectMatrix*’ tool from HiCExplorer.

#### Detection of mis-assemblies

Mis-assemblies are a common problem in *de novo* or published assemblies using Illumina and/or PacBio technologies and it is important to remove them; otherwise they introduce significant errors in Hi-C based assemblies. Mis-assemblies are readily spotted as discontinuous regions in Hi-C contact matrices that do not follow the power law decay with respect to genomic distance (Lieberman-Aiden et al., 2009). By plotting Hi-C contacts, those mis-assemblies can easily be detected by eye. For example, Figure S3E shows one *D. virilis* scaffold from the (Drosophila 12 Genomes Consortium et al., 2007) that contains a mis-assembly. HiCAssembler provides two methods to remove mis-assemblies:

##### Automatic method

Most mis-assemblies can be detected automatically when they occur between scaffolds located in different chromosomes or far away from each other in genomic distance. To detect mis-assemblies we use the TAD detection algorithm of HiCExplorer based on the TAD-separation score. Mis-assemblies can be identified as positions in the genome in which the adjacent downstream and upstream regions share significantly less contacts compared to the global average. Thus, the problem is similar to that of identifying TADs based on local minima of the TAD-separation score (Figure S3E). When running HiCAssembler, a cut off of the TAD-separation score can be given to split scaffolds. A downside of this method is that scaffolds can be erroneously split at strong TAD boundaries. However, during the Hi-C assembly process any wrongly identified splits are put together again, while mis-assemblies that are not separated introduce serious errors in the Hi-C assembly. In other words, false positives are not problematic for the Hi-C assembly, while false negatives are. Thus, it is preferable to set a threshold that removes all mis-assemblies at the expense of some false positives (TAD boundaries).

Un-split mis-assemblies can be detected by visual inspection of the HiCAssembler output which documents the Hi-C assembly process (Figure S3E).

##### Manual method

Some mis-assemblies cannot be removed automatically as they may appear similar to any other TAD boundary when they are not far away from their correct genomic location; or because they are close to the borders of scaffolds or in small scaffolds where it is not possible to accurately compute the TAD-separation score. These mis-assemblies can be detected by visual inspection of the automatically generated support images produced by HiCAssembler or, after the Hi-C assembly is finished and the resulting Hi-C contact matrix of the assembly shows unusual high contacts far away from the main diagonal. For these cases, it is possible to instruct HiCAssembler where to add splits using the ‘*--split_positions_file*’ parameter. To facilitate the determination of manual splits, HiCAssembler integrates a GUI tool called ‘*plotScaffoldsInteractively*’ that allows researchers to look and zoom into any single scaffold and to identify exact genomic position of desired split points.

#### Creation of initial path graph

To keep track of the Hi-C assembly process, HiCAssembler uses a path graph. In this type of graph, nodes can only be connected to at most two other nodes and cycles are not allowed. HiCAssembler creates a path graph in which each node is a bin of the corrected Hi-C contact matrix and scaffolds are represented by paths connecting their corresponding nodes. For example, if scaffold *S* contains bins with IDs {10, 11, 12, 13, 14} and the genomic position of 10…14 within *S* are consecutive, a path joining 10-11-12-13-14 is created. The implementation of paths in HiCAssembler prohibits connecting (i.e. adding edges) the internal nodes of the path as this will create a graph that is not a path. In case of example *S* those nodes are 11 to 13. In other words, once a path is created, new connections are only allowed when involving the first or last node.

HiCAssembler internally maintains two path graphs. Apart from the path graph joining bins of the Hi-C contact matrix, a second graph is constructed in which scaffolds are nodes (Figure S3B) and, as the Hi-C assembly progresses, paths of scaffolds are created in sync with larger paths connecting their corresponding bin paths.

#### Removal of small scaffolds and user-defined problematic scaffolds

In the next step, tiny scaffolds are removed from the graph path. Removed scaffolds are put aside and later put back into the Hi-C assembly. The length threshold to remove scaffolds is a parameter given by the user but in our analyses we have found that small scaffolds whose size is < 100 kb tend to introduce errors in the assembly. Smaller scaffolds share fewer contacts with other scaffolds and are thus less reliably ordered. Thus, using long scaffolds to build an initial Hi-C assembly results in better assemblies. Some scaffolds may still contain repetitive regions that appear as single. During Hi-C assembly, those scaffolds introduce ambiguities which can result in assembly errors and can be manually removed from the initial Hi-C assembly steps.

#### Iterative joining of high-confidence scaffold paths

HiCAssembler progresses by iteratively joining scaffolds to form larger and larger paths in each iteration until chromosome length assemblies are obtained (Figure S3A). During each iteration the individual steps taken are: 1. Merging of the initial matrix bins to create a smaller matrix. 2. Determination of a confidence cut off score. 3. Transformation of the merged matrix into a graph and computation of the maximum spanning tree. 4. Resolution of hubs and 5. Orientation and extension of scaffolds. In the following we will use the term Hi-C scaffold to refer to an already joined and oriented set of scaffolds.

The first iteration starts with pre-assembled contigs or scaffolds. The Hi-C scaffolds output from each iteration are the input for the next iteration.

##### 1. Merging initial matrix bins

During each iteration the Hi-C matrix is reduced by merging bins that belong to one Hi-C scaffold (Figure 1C). For each Hi-C scaffold, its internal bins are merged into parts that are about the size of the smallest Hi-C scaffold. Thus, some Hi-C scaffolds’ bins are all merged together into one new bin while other scaffolds may contain several new larger bins (Figure S3B second panel). Merging bins allows more Hi-C data to be aggregated per bin to gain confidence in the analysis. HiCAssembler uses fast algorithms that efficiently merge matrix bins. Because each new bin is the result of merging a different number of smaller bins, the new matrix is corrected using an optimized version of the iterative correction method (Imakaev et al., 2012).

##### 2. Determination of a contact cut off threshold to keep chromosomes separated

To estimate the number of contacts that are shared between scaffolds that are consecutive or separated by the average Hi-C scaffold length, we compute the median number of contacts between all parts of divided Hi-C scaffolds at all distances. These values are used to determine a cut-off threshold that will remove any contact between bins below this threshold. The cut-off threshold is set to the median number of contacts between Hi-C scaffolds that are separated by the length of one Hi-C scaffold. This will avoid joining scaffolds from distinct chromosomes, but also avoids joining Hi-C scaffolds that are separated from each other by at least the distance of the smallest Hi-C scaffold in each iteration. This step is different from strategies used by LACHESIS (Burton et al., 2013) and 3D-DNA (Dudchenko et al., 2017) to differentiate chromosomes: LACHESIS requires an initial clustering of scaffolds into a given number of groups while 3D-DNA uses a mis-assembly removal algorithm after the assembly process to separate the ’mega scaffold’ into chromosomes. In our opinion, this strategy makes the Hi-C assembly simpler by avoiding the initial clustering or the final separation of chromosomes.

##### 3. Transformation of the merged matrix into a graph and computation of the maximum spanning tree

The merged contact matrix is transformed into a weighted graph where each node is a Hi-C scaffold (or a part of a large Hi-C scaffold) and each edge weight is the corrected number of contacts shared by one pair of Hi-C scaffolds in the merged matrix (Figure S3B,C). The cut-off threshold from the previous step is applied to remove all edges whose weight is below the threshold (Figure S3 second panel). A maximum spanning tree (MST) is applied to this graph as in LACHESIS (Korbel and Lee, 2013) (Figure S3 third panel). The MST algorithm removes any cycles in the graph and leaves only edges with the highest weight. The graph before the MST algorithm and after the MST is saved in .graphml format. Those graphs can be visualized using for example Cytoscape (Shannon et al., 2003) and can be useful to identify problematic nodes that can afterwards be removed from the assembly using the ‘*--scaffolds_to_ignore*’ option.

##### 4. Resolution of hubs

After applying the MST, the resulting graph may contain nodes that are connected by more than two other nodes. We refer to these nodes as hubs. Otherwise, the MST graph mostly contains paths representing Hi-C scaffolds that can be joined to form larger Hi-C scaffolds. During the first iteration (before any scaffold has been attached to others) any branch of size one is pruned from the graph. These pruned nodes are put aside and integrated together with the smaller scaffolds after the Hi-C assembly of larger scaffolds finishes. Other hubs are resolved by leaving the top two edges with the highest weight and removing all other edges (Figure S3C last panel).

##### 5. Orientation and extension of scaffolds

After hub removal, the graph contains only paths in which each node is either a complete Hi-C scaffold or part of a divided large Hi-C scaffold (Figure 3B last panel). New connections between Hi-C scaffolds are now added. The following steps are carried out to resolve the orientation of scaffolds: *i*) identify all unmerged high-resolution bins that belong to each part of the Hi-C scaffolds (Figure 3D left). High-resolution bins correspond to bins in the original Hi-C matrix before step 1 of the iteration. *ii*) A small Hi-C matrix, containing only the selected bins is created. *iii*) Bins in the matrix are rearranged by keeping all bins that belong to one scaffold either in the same order (forward orientation) or in the inverted order (reverse orientation) (Figure S3D right). For each possible scaffold orientation the hic-score is computed using the following equation:

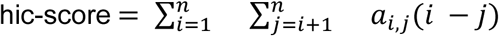

Here, *n* is the size of the small submatrix and *a_i,j_* is the value in the matrix for row *i* and column *j.* The orientation of scaffolds that results in the matrix that minimizes the hic-score is used to join and orient Hi-C scaffolds. Internally, the path graph of matrix bins and the path graph of scaffolds are updated.

In the hic-score function, values that are away from the main diagonal are multiplied by an increasingly higher number (*i - j*). Thus, only those Hi-C matrices having a high number of contacts close to the main diagonal will have low scores. This is expected because a correct Hi-C conformation is characterised by an approximate exponential decrease in the number of counts when moving away from the main diagonal of the matrix. This can be seen in the submatrices depicted in Figure S3D (right).

At the end of each iteration an image of the complete Hi-C matrix is saved. Because the hardware requirements grow quadratically with respect to the matrix size it is not practical to plot a matrix that has more than ∼4000 bins. Thus, the Hi-C matrix is reduced to at most 4000 bins if needed. These images are useful to detect problems with the assembly and to decide if adjustments are needed as for example manual splitting or removal of (erroneous parts of) scaffolds.

#### Addition of scaffolds that were not used yet

Once the iterative joining of Hi-C scaffolds ends, small scaffolds that were not used yet are put back into the Hi-C assembly. For this, paths of removed scaffolds are identified and inserted next to the Hi-C scaffold node with the highest number of contacts. In detail, first, a cut-off threshold is computed as in step 2, but this time the median of contacts for consecutive bins is used. Then, all scaffold bins are merged to form a smaller matrix as in step 1, the matrix is converted to a graph whose edges have weight = number of corrected contacts if the edge connects a removed scaffold. Otherwise, if the edge connects two Hi-C scaffold nodes already ordered and oriented, the edge weight = max(number of corrected contacts in the graph). In other words, all edges between scaffolds that were already joined in the iterative assembly have a maximum value. Afterwards, the cut-off is applied to remove low scoring edges. Then the MST is computed. Because all edges between Hi-C scaffolds already joined have the maximum value, none of those edges is removed in the MST computation. This creates a graph in which the removed scaffolds either form branches that are attached to a single Hi-C scaffold or are an independent tree. Next, we iterate over each branch and tree; if the branch/tree forms a path, the orientation of its scaffolds is determined as in step 5. If the branch is connected to a Hi-C scaffold, the branch is inserted into the Hi-C assembly scaffolds (corresponding to full-length chromosomes at this stage), otherwise the path is added to the Hi-C assembly as an unplaced scaffold.

#### Saving of scaffolds fasta file and liftover chain file

All Hi-C scaffolds are then saved as a fasta file. The header of the fasta file is composed by a unique id followed by a description of the scaffolds/contigs that were used their orientation. Between scaffolds a sequence of 2000 Ns is added. Additionally, a liftover chain file is saved. This file allows transferring annotation from the scaffolds to the Hi-C assembly.

### *In situ* Hi-C data processing

Paired-end reads were mapped and Hi-C matrices generated and corrected at restriction enzyme resolution as described above. We generated two replicates of stage 15-16 *D. melanogaster, D. virilis* and *D. busckii* embryos for further analysis and one replicate of 21-23 h old *D. busckii* embryos that was used for its genome assembly only. Hi-C matrices of replicates were merged using ‘*hicSumMatrices*’, matrix bins were merged using ‘*hicMergeMatrixBins*’ and matrices corrected afterwards using ‘*hicCorrectMatrix*’ tools from HiCExplorer v1.8.1. TAD boundaries were called using the ‘*hicFindTADs*’ tool from HiCExplorer with settings *’--minDepth 15000 --maxDepth 50000 --step 2000 --thresholdComparisons 0.01 --correctForMultipleTesting bonferronì*. In total we sequenced 45.7 M, 51.8 M and 64.5 M useful Hi-C reads from stage 15-16 *D. melanogaster, D. virilis* and *D. busckii* embryos, respectively and 26.8 M useful Hi-C reads from 21-23 h old *D. busckii* embryos (Table S2). For validation of our TAD calling, we used Hi-C datasets from Kc167 cells (Eagen et al., 2017) which we processed as described above.

First eigenvector (PCA1) corresponding to active (A) and inactive (B) compartments was computed using ‘hicPCA -noe1 --norm’ from HiCExplorer v2.2 after removal of heterochromatic chromosome ends. Corrected Hi-C matrices at restriction fragment resolution with 50 adjacent bins merged were used, resulting in matrices with a median bin size of about 25 kb. The correct orientation of PCA1, *i.e.* positive values corresponding to the active compartment (A) and negative values corresponding to the inactive compartment (B), was verified for each chromosome using female H4K16ac ChIP-seq data (this study).

### *D. busckii* Hi-C scaffolding

To perform *D. busckii* genome scaffolding using Hi-C data we used our 245 *de novo* contigs obtained by the Illumina and PacBio hybrid approach. For the assembly using HiCAssembler we used the following parameters: i)’*--min_scaffold_length 200000*’, to restrict the iterative scaffold assembly to scaffolds of 200 kb or bigger. Scaffolds smaller than 200 kb are added after the iterative correction. ii) ‘*--bin_size 10000*’, which sets the Hi-C bin size to 10 kb. This would be the size of high-resolution bins referred to in the algorithm description. iii) ‘*--misassembly_zscore_threshold −1.0*’ to control the threshold deciding if a TAD-separation score is strong enough to be considered a mis-assembly. iv) Additionally ‘*--scaffolds_to_ignore Backbone_81/13 Backbone_60/2 Backbone_59/2 Backbone_4/1 Backbone_53/13 Backbone_24/17 Backbone_88 Backbone_103/3*’ is used to ignore these contigs during the iterative assembly. Those scaffolds were ignored since they probably contain numerous repetitive regions (they appear as hubs in the maximum spanning tree algorithm, which can break the continuity of the assembly). We also set the number of iterations to 3 and defined a manual list of splits defined using ‘*plotScaffoldsInteractively*’.

Next we run whole genome alignments using NUCmer (NUCleotide MUMmer of mummer v4.0.0beta) with default parameters between the *D. busckii* fasta file produced by HiCAssembler and the fasta file for *D. melanogaster.* The chromosome names in the *D. busckii* assembly were set accordingly to the corresponding name in *D. melanogaster*.

### *D. virilis* Hi-C scaffolding

The *D. virilis* genome was sequenced and assembled into scaffolds as part of the Drosophila 12 Genomes Consortium (Drosophila 12 Genomes Consortium et al., 2007). These 13,415 dvir_caf1 scaffolds were downloaded from Ensembl (Assembly: GCA_000005245.1) (http://metazoa.ensembl.org/Drosophila_virilis/Info/Index). In the dvir_caf1 assembly, contigs were joined with a separation of variable number of stretches of “NNN”s in between. For the Hi-C assembly we split scaffolds that were separated with 10,000 or more Ns as we identified mis-assemblies associated with this scaffolds. For the Hi-C assembly we used the following parameters: ‘*--min_scaffold_length 100000 --bin_size 5000 --misassembly_zscore_threshold −1.0 --num_iterations 2*’. As for the *D. busckii* assembly, we used a whole genome alignment to the *D. melanogaster* genome to assign respective names to all chromosomes.

### *D. virilis* annotation liftover

We use the Hi-C scaffolding information to create a chain file to map the available annotation of *D. virilis* (dvir-all-r1.06.gtf) (ftp://ftp.flybase.net/releases/current/dvir_r1.06/) to the new Hi-C assembly using CrossMap v0.2.5 (Zhao et al., 2013).

### Synteny block detection

To identify synteny blocks (SBs) we use lastz (Harris, 2007) with the following parameters: ‘*--gfextend --nochain --gapped*’ which identifies local alignment blocks within respective chromosome arms obtained after extension (allowing for gaps) of seeds. We then chained blocks that are within 10 kb distance, that have the same orientation and that contained at least four lastz-defined blocks. Chained results that were smaller than 4 kb or completely overlapped a bigger synteny block were removed. The 10 kb merge distance was based on the longest intron length found in flies. Defining synteny block start and end sites as synteny breakpoints, we detect 3726 and 3252 breakpoints in the *D. melanogaster* vs. *D. virilis* comparison, respectively and 3340 and 2776 breakpoints in the *D. melanogaster* vs. *D. busckii* comparison, respectively.

### Overlaps between SB and TAD start and end sites

In order to shuffle the position of TADs and SBs along the genome while keeping the same region size distribution within the same chromosomes we used ‘*bedtools shuffle –noOverlapping-chrom -g chrom.size*’ with the adequate chromosome sizes attributed to the ‘*-g*’ parameter depending on the species. To calculate the overlap of SB and TAD start and end sites with SB breakpoints we extended TAD boundaries as well as SB start and end sites by 500 bp in 5’ and 3’ direction and calculated the fraction of overlap using ‘*bedtools intersect*’ and checked for significance using Fisher statistics ‘*bedtools fisher*’ with the ‘*-g*’ parameter according to the species.

### Calculation of Jaccard similarity index

The Jaccard similarity index was defined as the length of the intersection (in bp) divided by the length of the union of SBs and TADs (in bp):

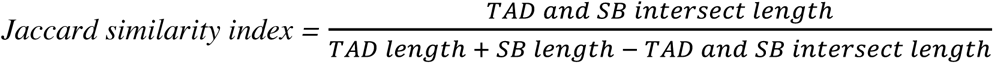

If one SB overlapped several TADs, we fused these TADs into one if the SB overlapped the adjacent TAD by at least 20 percent. This was done to account for several TADs keeping their linear order and being represented in one SB. Without this fusion of TADs, the Jaccard similarity index of one SB perfectly overlapping multiple TADs would result in a low Jaccard similarity index for each TAD, while the above mentioned method results in one fused TAD with a high Jaccard similarity index.

### Aligning TADs in between species using BLASTn

Sequences of *D. busckii* and *D. virilis* TADs were aligned to the sequences of *D. melanogaster* TADs by using BLASTn (blast+ v2.6.0, (Altschul et al., 1990; Camacho et al., 2009)). First, a database of *D. melanogaster* TAD sequences was made using *’makeblastdb -dbtype nucl’.* Then, the sequences of *D. virilis* and *D. busckii* TADs were compared to this database and the best hit was retrieved using ‘*blastn -outfmt 6 -evalue 1000 -max_target_seqs 1’*. Using this method, a BLASTn bitscore was assigned to each *D. virilis* or *D. busckii* TAD compared to *D. melanogaster* TADs, reflecting how well a given TAD in one species is conserved in the other species.

### Definition of conserved TADs

TADs in *D. melanogaster* with a Jaccard similarity index or bitscore (see paragraph before) above the median from the comparison with *D. busckii* or *D. virilis* were overlapped using ‘*bedtools intersect -f 0.8 -r*’. The intersect of both comparisons was overlapped and resulted in the definition of 175 conserved TADs. These 175 conserved TADs cover 11 Mb of the *D. melanogaster* genome (see Figure 4A).

### Characterization of conserved TADs

Conserved TADs were compared with unconserved TADs and random regions. Unconserved TADs were defined as TADs with a similar length distribution as the conserved TADs (by selecting TADs of sizes between the 0.05th and 0.95th quantile of conserved TADs) and excluding TADs with high Jaccard and bitscores used in the intersect for the definition of conserved TADs. We selecting TADs with a similar length distribution because conserved TADs are larger than average TADs (see Figure S6A). Random regions were defined as genomic regions with the same length distribution inside the same chromosomes as conserved TADs and are not corresponding to TADs but random genomic regions by using ‘*bedtools shuffle -noOverlapping -chrom*’.

Conserved TADs in *D. melanogaster* were overlapped with all genes (Ensembl Genes 92, fruit fly genes (BDGP6)) and the number of overlapping genes per kb was calculated. The distribution of the gene length of overlapping genes is plotted in Figure S6B as a control.

Enrichment of NSL3 at the boundaries of conserved TADs was shown using NSL3 ChIP-seq data from *D. melanogaster* S2 cells (Lam et al., 2012). Fastq files were mapped using Bowtie2 (Langmead and Salzberg, 2012) with default parameters. Log2fold ratio over the input sample was calculated using bamCompare (deeptools v3.0.2) with with settings ‘*-bs 5 -ignore chrX*’.

Overlap of conserved TADs with chromatin states was calculated using chromatin states reported in *D. melanogaster* Kc cells with the accession number GSE22069 (Filion et al., 2010). The bed file was lifted over from Dm3 to Dm6 using CrossMap (Zhao et al., 2013). The significance of the difference to unconserved TADs and random regions was tested using Wilcoxon rank-sum tests (Figure S6C).

Enrichment of H3K4me3, H3K36me3, H3K27me3 and Hp1 alpha in conserved TADs was shown using ChIP-seq data of 14-16 h old *D. melanogaster* embryos from the modENCODE Project (modEncode accession 5096, 4950, 3955 and 3956) (Celniker et al., 2009). Data processing included sequencing quality and adaptor trimming of single-end reads using trim_galore_v0.4.5 (<u>bioinformatics.babraham.ac.uk</u>) with setting ‘*-q 5*’, mapping using Bowtie2 v2.3.4.1 (Langmead and Salzberg, 2012). The coverage of mapped reads was calculated using deeptools v3.0.2 (Ramírez et al., 2016) with settings ‘*-bs 25*’. Log2fold ratio over the input sample was calculated using bamCompare (deeptools v3.0.2) with settings ‘*-bs 25*’.

### H4K16ac ChIP-seq experiments

ChIP was performed as described in (Valsecchi et al., 2018). Briefly, male and female 3rd instar larvae were separated and carcasses dissected followed by fixation with 0.2 % formaldehyde for 15 min. Extraction was performed using a pestle homogenizer in 0.2 mL buffer G1 (15 mM HEPES (pH 7.4), 20 mM KCl, 10 mM MgCl_2_, 0.5 % Tween-20, 20 % glycerol with protease inhibitors) followed by a sucrose cushion and a subsequent wash step with 10 mM Tris (pH 8.0), 5 mM MgCl_2_, 1 mM CaCl_2_, 10 mM NaCl, 0.1 % IGEPAL CA-630, 0.25 M sucrose. Pelleted nuclei were resuspended in 20 mM Tris (pH 8.0), 5 mM MgCl_2_, 1 mM CaCl_2_, 10 mM NaCl, 1 % Triton-X100, 0.25 M sucrose for digestion with Micrococcal Nuclease (M0247, New England Biolabs NEB). After 15 min digestion at RT, 33 μL of 10x high salt buffer were added (200 mM EDTA, 4 M NaCl) before treatment in a Bioruptor Pico (10 cycles, 30 sec ON / OFF). The extracts were clarified by centrifugation for 5 min at 12’000 x g. ChIP was performed on this chromatin extract using 1 µL of Anti-acetyl-Histone H4 (Lys16) Antibody (Merck Milipore, 07-329) or 0.5 µL of Histone H3 antibody mAb MABI 0301 (Active Motif, 39763). Immunoprecipitated chromatin was captured using Dynabeads Protein A (Thermo Fisher, 10001D). After reverse crosslinking in 1x TE at 65°C for 16 hours, RNaseA treatment and proteinase K-treatment, the DNA was phenol-chloroform extracted followed by ethanol precipitation. Library preparation for sequencing was performed using a NEBNext® Ultra™ II DNA Library Prep Kit for Illumina (E7645, New England Biolabs NEB). Samples were sequenced paired-end, with a read length of 75 bp, on a Illumina HiSeq 3000.

### H4K16ac ChIP-seq data processing

We analysed our generated H4K16ac ChIP-seq data from *D. virilis* and *D. busckii* (see paragraph before) as well as our already published H4K16ac ChIP-seq data from D. melanogaster (GSE109901). Data processing included sequencing quality and adaptor trimming of paired-end reads using trim_galore v0.4 (<u>bioinformatics.babraham.ac.uk</u>), mapping individual replicates using bwa v0.7.12 (<u>arXiv:1303.3997v2</u>) followed by sorting and indexing of bam files using samtools-1.2 (Li et al., 2009). The coverage of mapped reads was calculated using deeptools v2.0.1 (Ramírez et al., 2016) with settings ‘*-bs 10 --minMappingQuality 2 --normalizeTo1x {EFFECTIVE_GENOME_SIZE}*’. We defined the effective genome size as the total genome size minus the number of Ns which is 117 Mb, 188 Mb and 120 Mb for *D. busckii, D. virilis* and *D. melanogaster*, respectively. Log2fold ratios of merged replicates over the input sample was calculated using bamCompare (deeptools v2.0.1) with settings ‘*-bs 10 --scaleFactorsMethod SES*’.

### Repeat modelling and repeat masking of genomes

We first performed *de novo* repeat discovery using RepeatModeler (v1.0.10) (repeatmasker.org) with default settings. Afterwards, we combined the *de novo* discovered repeats with 2385 *Drosophila* and ancestral repeats from Repbase (Bao et al., 2015). Then we run RepeatMasker v4.0.5 (repeatmasker.org) with this combined library of repeats and default settings. All three genomes were treated the same.

### Analysis of TAD boundary motif enrichment

We used a list of boundary motifs (Ramírez et al., 2018) that we have described in *D. melanogaster* to analyze and compare their enrichments at TAD boundaries in all three species using AME (McLeay and Bailey, 2010) from the MEME suite v.5.0.2. Our list contains the following motifs: Beaf-32, CTCF, GAATAGAAA, GAF, Ibf, Ohler-1, Ohler-5, Ohler-6, Ohler-8, Pita, Su(Hw), Su(Hw)_short, ZIPIC, Zw5 and Ohler-8_dreme. We extracted 500 bp around all TAD boundaries in repeat-masked genomes (see paragraph before), removed sequences with more than 100 Ns and build a 2nd order model of TAD boundary sequences as the background model. We used shuffled input sequences a control, average odds score as the sequence scoring method and one-tailed Wilcoxon rank-sum test as the motif enrichment test.

To compare motifs that we have described to be enriched at promoter (Ohler-1, Beaf-32, Ohler-6, ZIPIC and Ohler-8) or non-promoter (CTCF, Su(Hw), Ibf) boundaries (Ramírez et al., 2018) in conserved TAD boundaries vs. unconserved TAD boundaries, we analyzed enrichment of these motifs using AME.

### Analysis of Beaf-32 ChIP-seq

Fastq files for Beaf-32 ChIP-seq and input from <u>GSM762845</u> (Van Bortle et al., 2014) were downloaded and aligned to the Dm3 assembly using Bowtie2 (Langmead and Salzberg, 2012). MACS2 (Zhang et al., 2008) was used to identify peaks. bamCompare and bamCoverage from deepTools2 (Ramírez et al., 2016) were used to create normalized coverage bigwig files. The processed files were lifted over from Dm3 to Dm6 using CrossMap (Zhao et al., 2013).

### Polytene chromosome spreads

Polytene chromosomes from separated male and female third-instar larvae in all three *Drosophila* species were prepared as previously described (Zink and Paro, 1995). Briefly, fixed and blocked polytene chromosome spreads were incubated with a home-made primary anti-MOF antibody (Mendjan et al., 2006) (1:400 in goat serum). The secondary antibody (Alexa 488 goat anti-rabbit, A-11034 from Thermo Fisher Scientific) was used in a 1:500 dilution together with Hoechst as well in a 1:500 dilution. Images were obtained with a Zeiss Elyra system (Carl Zeiss Microscopy) and processed using Fiji (Schindelin et al., 2012).

### roX2 ChIRP-seq data processing

ChIRP-seq data from *D. melanogaster, D. virilis* and *D. busckii* was downloaded from GEO (GSE69208) (Quinn et al., 2016). Data processing included sequencing quality and adaptor trimming of single-end reads using trim_galore v0.4 (<u>bioinformatics.babraham.ac.uk</u>), mapping individual replicates (odd and even) using bwa v0.7.12 (<u>arXiv:1303.3997v2</u>) followed by sorting and indexing of bam files using samtools-1.2 (Li et al., 2009). The coverage of mapped reads was calculated using deeptools v2.0.1 (Ramírez et al., 2016) with settings ‘*-bs 10 --normalizeTo1x {EFFECTIVE_GENOME_SIZE}*’. Log2fold ratios of merged replicates (odd and even) over the input sample was calculated using bamCompare (deeptools v2.0.1) with settings ‘*-bs 10 --scaleFactorsMethod SES*’. Peak calling was done using MACS2 v2.1.1.20160309 (Zhang et al., 2008) with settings ‘*callpeak -f BAM --qvalue 0.01 -g {effective genome size}*’. ChIRP samples from different species were sequenced to a different depth which affects the -log10(q-value) of peaks called using MACS2. Due to this reason and to have a comparable number of roX2 peaks for further analysis, we used the ‘*--normalizeTo1x*’ coverage files to select 250 high-confidence roX2 peaks per species (referred to as high-affinity sites (HAS)). The criteria to select those peaks were: a minimum normalized coverage of 50 (meaning 50 times the enrichment over background), the highest -log10(q-value) and the peak needs to be present in the two replicates. The peak summit was identified as the location of the highest coverage value.

### Aggregated Hi-C contacts

We used hicAggregateContacts from HicExplorer with corrected Hi-C matrices and settings ‘*--vMin 1 --vMax 2 --range 300000:1000000 --numberOfBins 30 --chromosomes X --avgType mean --transform obs/exp*’ to plot aggregated Hi-C contacts of high-confidence roX2 binding sites (HAS) on the X chromosome in matrices with 3 bins merged (∼1.7 kb bins size). Out of the selected 250 HAS, we found 246 in *D. melanogaster*, 247 in *D. virilis* and 213 in *D. busckii* to be located on the X chromosome. We choose the respective number of random regions on the X chromosome (’*shuffleBed*’ from bedtools2) for comparison to random aggregated Hi-C contacts.

Enriched Hi-C contacts between the respective number of TAD boundaries on the X chromosome of the lowest z-score were visualized using aggregate plots as described above for HAS. Distances of HAS to the closest TAD boundary were added on the opposite side of the TAD boundary to get “mirrored” HAS. Aggregated Hi-C contacts were plotted as described above. Gene expression (normalized counts) analyses was performed using library size normalized RNA-seq counts from 14-20 h aged embryos from modENCODE datasets obtained from (Ramírez et al., 2018) and also available on the chorogenome web server (http://chorogenome.ie-freiburg.mpg.de/). Overlap of HAS with 5’ UTR, exon, intron, 3’ UTR and intergenic regions was calculated using segtools-overlap (Buske et al., 2011).

### HAS to HAS Hi-C contact visualization

Figure 5B shows total HAS to HAS obs/exp Hi-C contacts (red arcs) displayed using pyGenomeTracks v2.0 (https://github.com/deeptools/pyGenomeTracks). The total HAS to HAS contacts were retrieved by converting Hi-C matrices at restriction enzyme sites resolution with 3 adjacent bins merged using ‘*hicMergeMatrixBins*’ into obs/exp matrices with ‘*hicTransform*’ that were then exported in GInteractions format using ‘*hicExport*’ from HiCExplorer v2.1.4. Then anchors comprised in GInteractions files were overlapped with the HAS defined in each species using the ‘*InteractionSet*’ R package (Lun et al., 2016).

### Analysis of phenotypic classes of alleles

The essentiality of genes overlapping conserved TADs (Figure 4I) was assessed in *D. melanogaster* using the Flybase automated gene summaries. The genes were filtered for genes with phenotypic annotation in *D. melanogaster* (i.e. 21 671 FBgn IDs). The analysis consists of calculating the fraction of genes displaying one of the four groups of phenotypic classes aggregated from the 184 different terms found in the automated gene summaries: the “Lethal” class corresponds to genes with “; lethal” or “lethal -” annotation, the “Increased mortality” class corresponds to genes with “increased mortality” annotation, the “Some die” class corresponds to genes with “some die during” annotation, and the “Viable” class corresponds to genes with “viable” annotation. Genes can have multiple annotated phenotypic classes as different alleles can have different phenotypic effects that can be lethal, partially lethal or viable, thus the fraction displayed in Figure 4I do not sum to 1.

### Gene Ontology enrichment of Biological Processes

Gene Ontology enrichment of Biological Processes (GOBP) was computed using DAVID with default parameters (Huang et al., 2009a, 2009b). The list of enriched GOBP was processed by REVIGO for plotting and to further filter the terms using a small (0.5) allowed similarity (Supek et al., 2011).

### Statistics

All statistical tests are reported in the Figure legends. All boxplots show interquartile ranges (IQR, 0.25th to 0.75th quartile (Q_1_-Q_3_)), whiskers represent Q_1_- 1.5*IQR (bottom), Q_3_- 1.5*IQR (top) and notches represent the *median* 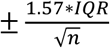.

