## Supplemental figures and tables for "Hi-C guided assemblies reveal conserved regulatory topologies on X and autosomes despite extensive genome shuffling"

#### Supplemental Information

This file contains:

- 7 Supplemental Figures and Figure Legends
- 3 Supplemental Tables and Table Legends
- Supplemental References

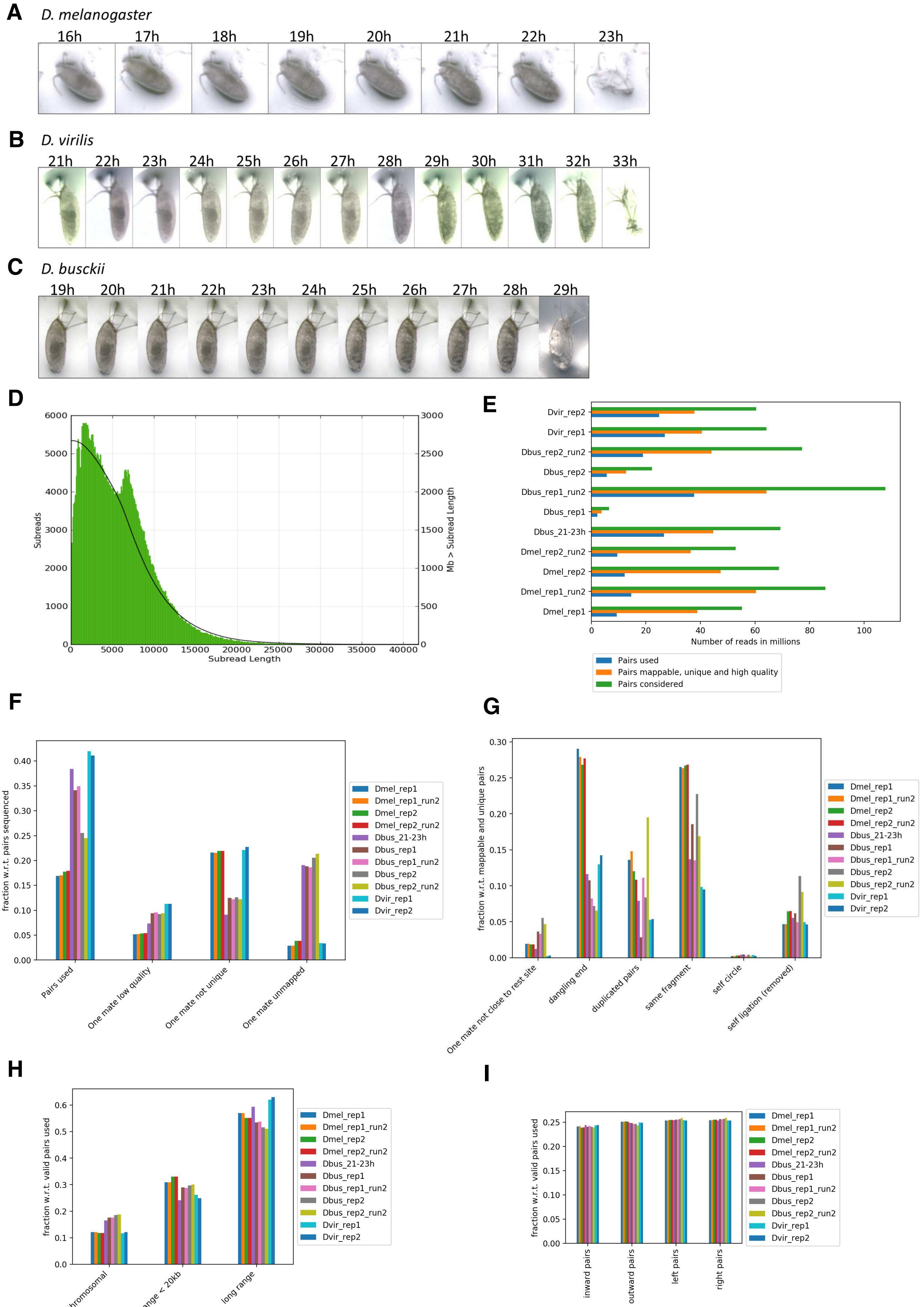

A
D. melanogaster
16h 17h 18h 19h 20h 21h 22h 23h
B
D. virilis
21h 22h 23h 24h 25h 26h 27h 28h 29h 30h 31h 32h 33h
C
D. busckii
19h 20h 21h 22h 23h 24h 25h 26h 27h 28h 29h
D
Subreads
Mb > Subread Length
Subread Length
E
Dvir_rep2
Dvir_rep1
Dbus_rep2_run2
Dbus_rep2
Dbus_rep1_run2
Dbus_rep1
Dbus_21-23h
Dmel_rep2_run2
Dmel_rep2
Dmel_rep1_run2
Dmel_rep1
Number of reads in millions
Pairs used
Pairs mappable, unique and high quality
Pairs considered
F
fraction w.r.t. pairs sequenced
Pairs used
One mate low quality
One mate not unique
One mate unmapped
Dmel_rep1
Dmel_rep1_run2
Dmel_rep2
Dmel_rep2_run2
Dbus_21-23h
Dbus_rep1
Dbus_rep1_run2
Dbus_rep2
Dbus_rep2_run2
Dvir_rep1
Dvir_rep2
G
fraction w.r.t. mappable and unique pairs
One mate not close to rest site
dangling end
duplicated pairs
same fragment
self circle
self ligation (removed)
H
fraction w.r.t. valid pairs used
inter chromosomal
short range < 20kb
long range
I
fraction w.r.t. valid pairs used
inward pairs
outward pairs
left pairs
right pairs

**Figure S1**

**Figure S1, related to Figure 1. Generated PacBio and Hi-C datasets.**
**(A,B,C)** Developing *D. melanogaster* **(A)**, *D. virilis* **(B)** and *D. busckii* **(C)** embryo from stage 15-16 until hatching. The embryos were kept at 25°C and imaged every hour. The midgut is characteristically visible at stage 15 as a dark spot in the center of the embryo. The tracheal tree becomes visible at stage 17, just before hatching.
**(D)** Histogram of the PacBio subread length. The mean subread length is 5.7 kb.
**(E)** Number of sequenced (green) mappable, unique and high quality (orange) as well as valid used (blue) Hi-C reads per sequenced sample. If sequencing of one sample was split into two runs, the second run is indicated with 'run2'. Exact numbers of sequenced and valid reads are available in Table S2.
**(F)** Fraction of used, unmappable and non-unique read pairs with respect to the sequenced read pairs.
**(G)** Fraction of read pairs (with respect to mappable and unique reads) that were discarded when building the Hi-C matrix.
**(H)** Fraction of inter-chromosomal, short-range (< 20 kb) and long-range Hi-C contacts with respect to the valid read pairs used.
**(I)** Fraction of inward, outward, left and right read pairs with respect to the valid read pairs used.

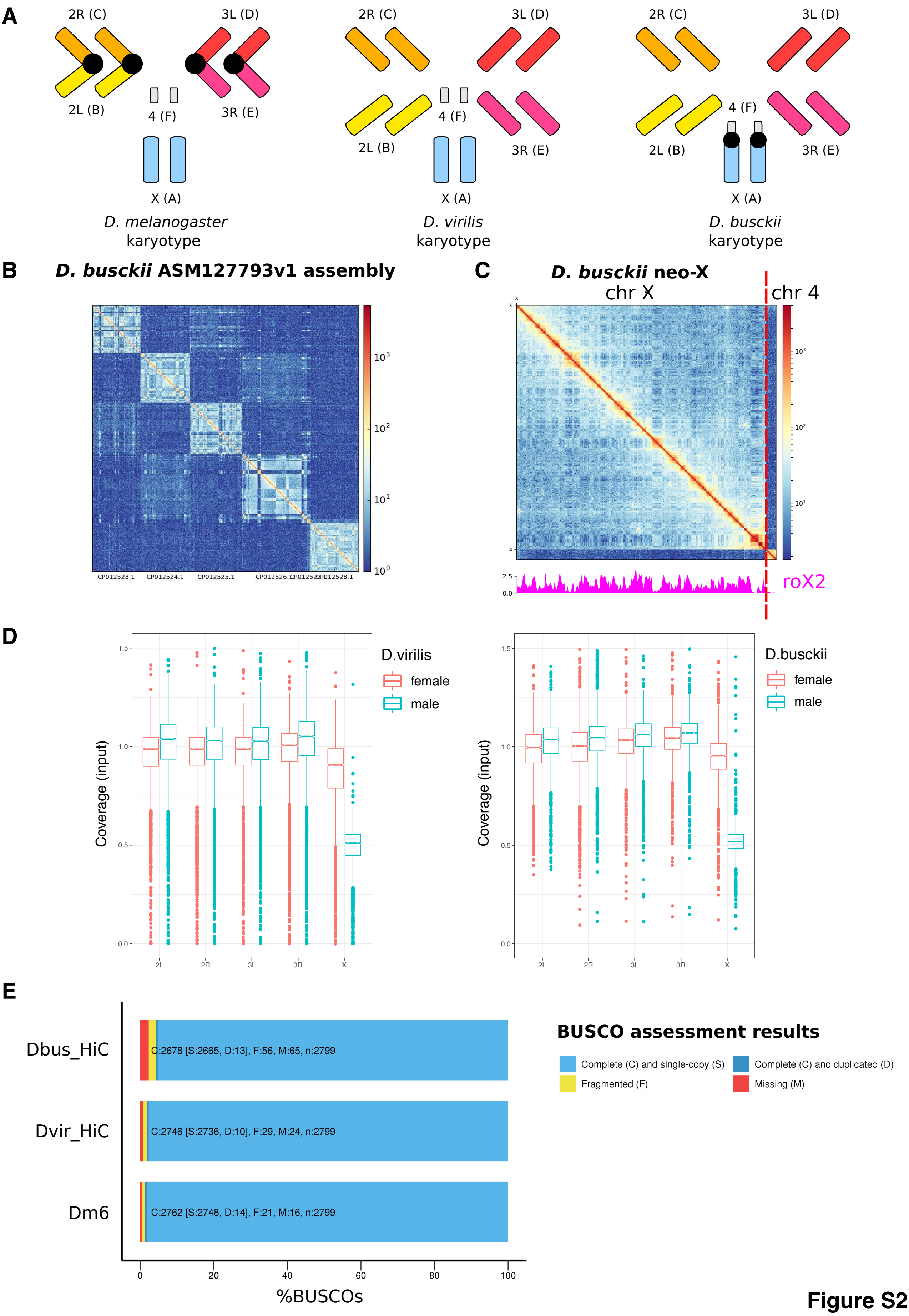
A
2R (C)
3L (D)
2L (B)
4 (F)
3R (E)
X (A)
D. melanogaster karyotype
D. virilis karyotype
D. busckii karyotype
B
D. busckii ASM127793v1 assembly
CP012523.1
CP012524.1
CP012525.1
CP012526.1
CP012528.1
C
D. busckii neo-X
chr X
chr 4
roX2
D
D.virilis
female
male
Coverage (input)
2L
2R
3L
3R
X
D.busckii
E
Dbus_HiC
Dvir_HiC
Dm6
C:2678 [S:2665, D:13], F:56, M:65, n:2799
C:2746 [S:2736, D:10], F:29, M:24, n:2799
C:2762 [S:2748, D:14], F:21, M:16, n:2799
%BUSCOs
BUSCO assessment results
Complete (C) and single-copy (S)
Complete (C) and duplicated (D)
Fragmented (F)
Missing (M)

**Figure S2**

**Figure S2, related to Figure 1. Hi-C assembly quality control.**

**(A)** Female karyotypes of *D. melanogaster, D. virilis* and *D. busckii.* Chromosomal arms X, 2L, 2R, 3L, 3R and 4 corresponding to Muller elements A, B, C, D, E and F are shown as colored boxes. Fusions of chromosome arms are indicated with black circles.

**(B)** The *D. busckii* ASM127793v1 assembly (current RefSeq representative genome) was generated by aligning assembled scaffolds to the *D. melanogaster* genome to generate pseudo-chromosomal sequences (Zhou and Bachtrog, 2015). The equivocated order of scaffolds is visible as abruptly changing Hi-C contacts within and outside of chromosome arms.

**(C)** roX2 ChIRP-seq data on the neo-X chromosome of *D. busckii*. In *D. busckii,* chromosome X (chr X) and small heterochromatic chromosome 4 (chr 4) are fused into one neo-X chromosome (Krivshenko, 1959, 1955). Zhou and Bachtrog showed that Muller element F (chr 4) is not enriched for H4K16ac despite its fusion to the X chromosome (Zhou and Bachtrog, 2015). Analysis of ChIRP-seq data of roX2 (Quinn et al., 2016) does not show enrichment of roX2 on chromosome 4 compared to the X chromosome as expected. The border between the X chromosome and chromosome 4 is highlighted with a red dashed line.

**(D)** Coverage of mapped reads of male and female ChIP-seq input sample which was normalized to 1x (see methods) per 10 kb bin. The coverage of the male X chromosome in approximately 0.5 compared to 1 in females which shows that it is present in a single copy in males and two copies in females.

**(E)** The *Diptera* (odb9) dataset containing 2799 Benchmarking Universal Single-Copy Orthologs (BUSCOs) (Waterhouse et al., 2017) was used to measure the completeness of our new genome assemblies (D_busckii_HiC and D_virilis_HiC) compared to the Dm6 genome. Reported are complete (C) and single-copy (S, light blue) or duplicated (D, dark blue) as well as fragmented (F, yellow) and missing (M, red) BUSCOs.

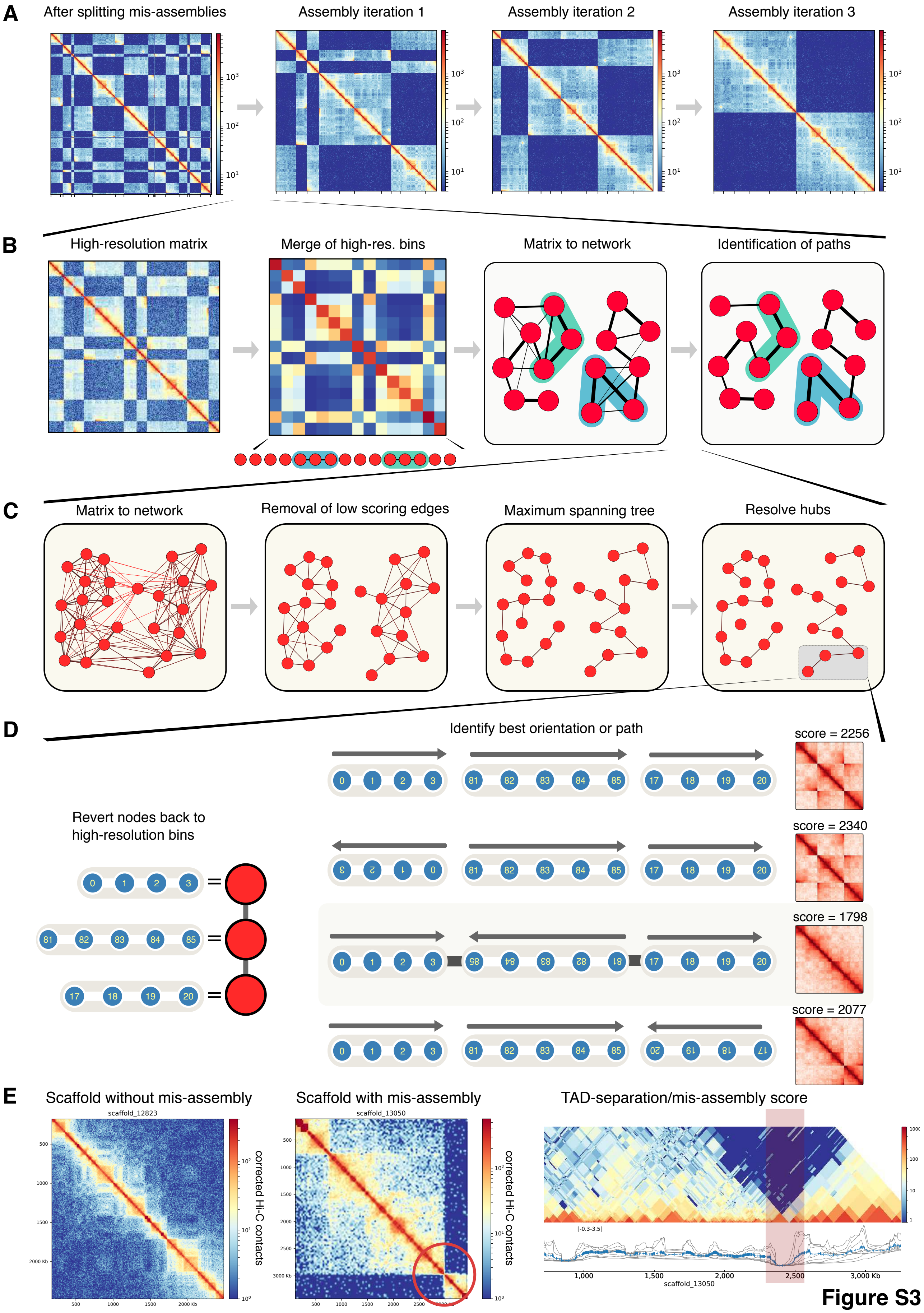
A
After splitting mis-assemblies
Assembly iteration 1
Assembly iteration 2
Assembly iteration 3
B
High-resolution matrix
Merge of high-res. bins
Matrix to network
Identification of paths
C
Matrix to network
Removal of low scoring edges
Maximum spanning tree
Resolve hubs
D
Identify best orientation or path
Revert nodes back to high-resolution bins
score = 2256
score = 2340
score = 1798
score = 2077
E
Scaffold without mis-assembly
Scaffold with mis-assembly
TAD-separation/mis-assembly score
scaffold_12823
scaffold_13050
corrected Hi-C contacts

**Figure S3**

**Figure S3, related to Figure 1. HiCAssembler.**

**(A)** Example Hi-C matrix depicting the iterative progression of the Hi-C assembly strategy as in (Dudchenko et al., 2017). First, the Hi-C data is mapped to scaffolds, a Hi-C matrix of restriction fragment bins is created and corrected. Then, the original scaffolds are split if they contain mis-assemblies and small scaffolds are removed (fist panel). In each iteration of the Hi-C assembly algorithm, scaffolds are joined and oriented to form what we call Hi-C scaffolds. After each iteration the Hi-C scaffolds become bigger until chromosome-length assemblies are obtained as shown in the last panel were two separated blocks remain. Afterwards, the small scaffolds that were initially removed are inserted into the Hi-C scaffolds. **(B)** Merging of high-resolution bins and determination of paths. In each iteration, all high resolution bins of the Hi-C matrix belonging to one of the Hi-C scaffolds are merged together (during the first iteration, Hi-C scaffolds are equal to the initial scaffolds). In the second panel (from left to right), each of the circles represent one Hi-C-scaffold whose high-resolution bins were merged. The new matrix is corrected to eliminate differences caused by merging Hi-C scaffolds of different length. Also, larger scaffolds are divided before merging them into equally long parts whose size is about the size of the smaller Hi-C scaffold. In the example shown, two large Hi-C scaffolds are divided into three parts; this is represented by the three connected circles with blue and green backgrounds. Using the divided scaffolds, the average number of corrected contacts between each part is computed to identify a cut-off threshold of the average number of counts between parts separated by one part. Next, the matrix is converted into a weighted graph to identify all Hi-C scaffolds that should be joined. In this graph, each node is a Hi-C scaffold (or a part of a Hi-C scaffold) and each edge's weight is the corrected number of contacts shared by any pair of Hi-C scaffolds. In the two rightmost panels, the Hi-C scaffolds that were divided are shown with the blue and green background and the edge weight is proportional to the line thickness.

**(C)** Example usage of a larger network. For simplicity, subdivided Hi-C scaffolds are not shown. After the merged Hi-C matrix is converted into a weighted graph, all edges having a weight below the cut-off threshold are removed. This divides the network into connected components representing groups of Hi-C scaffolds that are at most the length of one Hi-C scaffold appart in genomic distance. Then, the maximum spanning tree is computed and hubs are resolved by either pruning single nodes or by removing the weakest edges. The resulting graph contains the order in which the Hi-C scaffolds should be joined.

**(D)** Once the ordering has been determined, the orientation of all parts is decided. For this, we use the high-resolution bins that form each Hi-C scaffold and compute their hic-score (see methods) in different orientations. The set of orientations that minimizes the hic-score is used to join the Hi-C scaffolds into a larger Hi-C scaffolds. In the figure, an example of a path of three Hi-C scaffolds is shown together with its respective high resolution bins (left). Four possible combinations of orientations are shown. For each case, the high-resolution submatrix (in log scale) containing only the bins of the three hic-scaffolds is shown on the right together with the hic-score. The third combination of orientations is the correct one as this produces the typical Hi-C matrix.

**(E)** Depicted are two scaffolds of the Dvir_caf1 assembly, one without a mis-assembly (left) and one with a mis-assembly (middle). Mis-assemblies (red circles) are readily spotted as discontinuous regions in the Hi-C contact matrix that do not follow the power law decay with respect to genomic distance. Mis-assemblies can automatically be identified as positions that share significantly less contacts compared to the global average. This can be detected by HiCAssembler as minima in the TAD-separation/mis-assembly score (right).

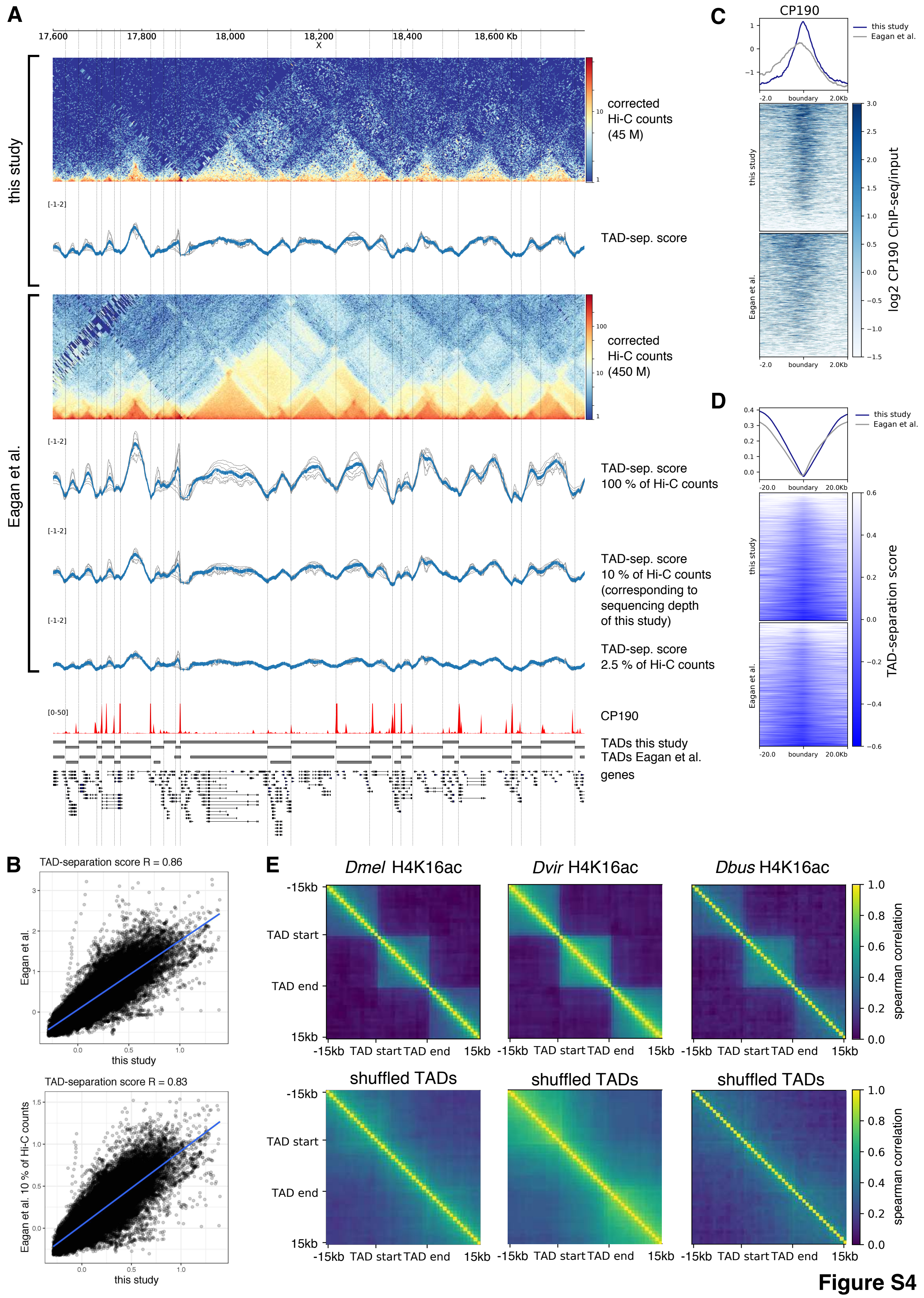
A
17,600
17,800
18,000
18,200
X
18,400
18,600 Kb
this study
corrected
Hi-C counts
(45 M)
TAD-sep. score
Eagan et al.
corrected
Hi-C counts
(450 M)
TAD-sep. score
100 % of Hi-C counts
TAD-sep. score
10 % of Hi-C counts
(corresponding to
sequencing depth
of this study)
TAD-sep. score
2.5 % of Hi-C counts
CP190
TADs this study
TADs Eagan et al.
genes
B
TAD-separation score R = 0.86
Eagan et al.
this study
TAD-separation score R = 0.83
Eagan et al. 10 % of Hi-C counts
this study
C
CP190
this study
Eagan et al.
boundary
log2 CP190 ChIP-seq/input
D
this study
Eagan et al.
boundary
TAD-separation score
E
Dmel H4K16ac
Dvir H4K16ac
Dbus H4K16ac
shuffled TADs
-15kb
TAD start
TAD end
15kb
spearman correlation

Figure S4

**Figure S4, related to Figure 2. TAD calling validation.**

**(A)** Example region on chromosome X showing a Hi-C contact matrix and TAD-separation score (TAD-sep. score) in *D. melanogaster* embryos (this study) and in a 10-fold deeper sequenced dataset from Kc167 cells (Eagen et al., 2017). The TAD-sep. score calculated using HiCExplorer is based on a z-score transformation of the Hi-C matrix as described in (Ramírez et al., 2018). TAD boundaries correspond to local minima of this score. TAD boundaries called using the TAD-sep. score in this study are shown as vertical lines. The TAD-sep. score calculated using HiCExplorer is robust to sequencing depth differences as shown by downsampling to 10 % (approximately corresponding to the sequencing depth used in this study) and 2.5 % of the Eagan et al. dataset. The following tracks show normalized ChIP-seq coverage for the common insulator protein cofactor CP190 (Li et al., 2015) (GSM1535980) as well as TADs called in this study using HiCExplorer (n= 2209) and TADs called by Eagan et al. using the Arrowhead algorithm (n=2127).

**(B)** Spearman correlation of the TAD-sep. score from this study (45M Hi-C contacts) and Eagan et al. (450M Hi-C contacts) as well as from this study and downsampled dataset from Eagan et al.

**(C)** Comparison of TAD boundaries called in this study and TAD boundaries called by (Eagen et al., 2017) with respect to CP190 (Li et al., 2015). Our boundaries overlap more frequently with CP190.

**(D)** Similar to (C), but comparing TAD-sep. score.

**(E)** Correlation of the histone mark H4K16ac (female 3rd instar larvae, this study) within and outside TADs (+15 kb flanking regions) as well as shuffled TADs as a control in *D. melanogaster, D. virilis* and *D. busckii*. Each pixel in the matrix represent the Spearman correlation of the histone mark in and outside of all TADs.

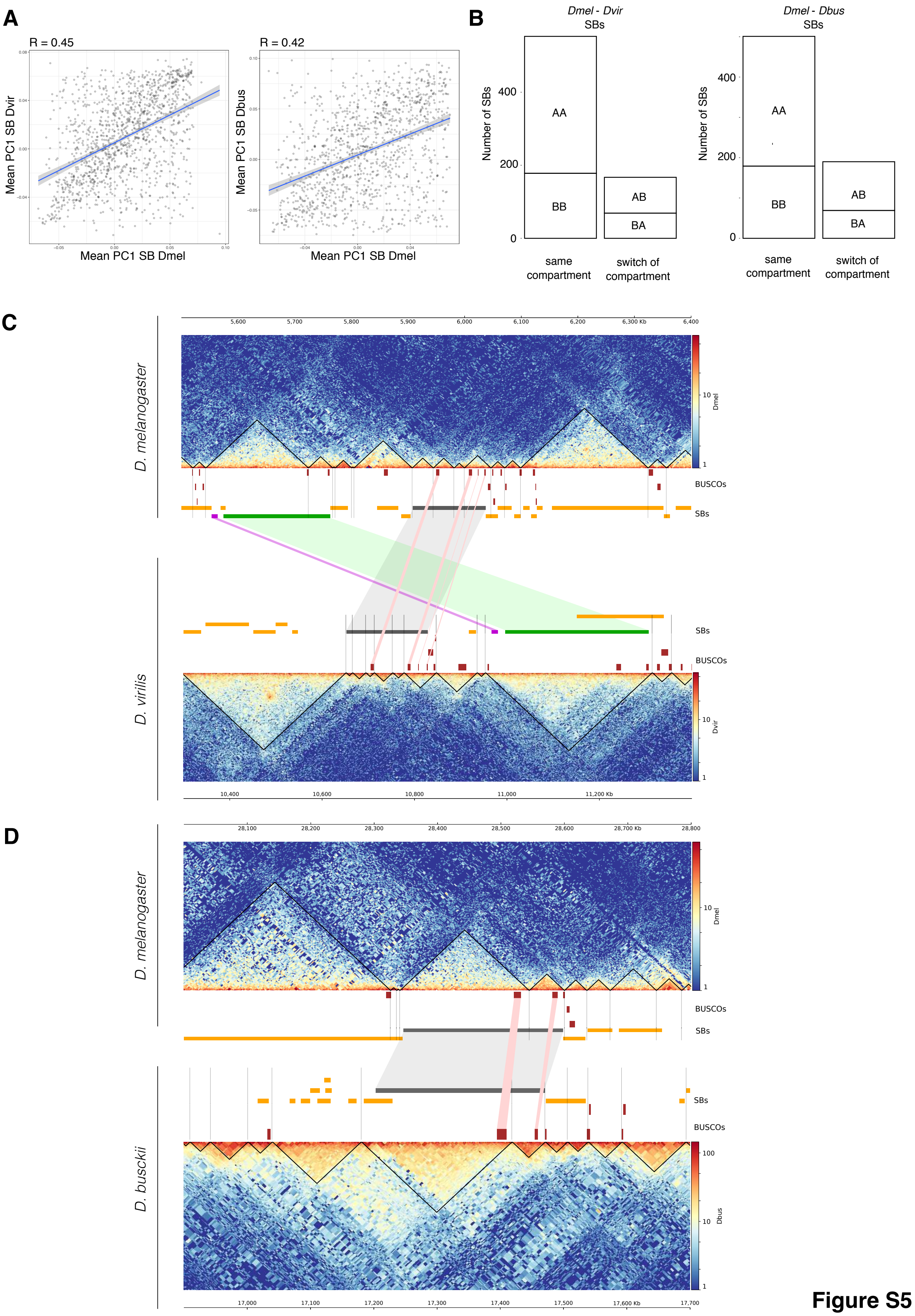
A
R = 0.45
Mean PC1 SB Dvir
Mean PC1 SB Dmel
R = 0.42
Mean PC1 SB Dbus
Mean PC1 SB Dmel
B
Dmel - Dvir
SBs
Number of SBs
AA
BB
AB
BA
same compartment
switch of compartment
Dmel - Dbus
SBs
C
D. melanogaster
BUSCOs
SBs
D. virilis
D
D. melanogaster
D. busckii

**Figure S5**

**Figure S5, related to Figure 2. Examples of conserved SBs between *D. melanogaster* and *D. virilis* or *D. busckii.***

**(A)** Pearson correlation of mean PCA1 per SB between *D. melanogaster* (DM) and *D. virilis* (DV) or *D. melanogaster* (DM) and *D. busckii* (DB) after removal of outliers (-0.1 < PCA1 < 0.1).

(B) Number of *Dmel- Dvir* or *Dmel- Dbus* SBs that lie completely in the A or B compartment in both species and stay in the same compartment (AA and BB) or switch compartment (AB and BA) between both species.

**(C)** Hi-C contact heatmaps including called TADs (black triangles), BUSCOs (red boxes) and SBs (orange boxes) in *D. melanogaster* and *D. virilis.* Correspondence between conserved SBs (gray, green and magenta boxes) is indicated with connections between the SBs in both species. Corresponding BUSCOs are indicated with connections between them.

**(D)** Comparison of a 800 kb region in between *D. melanogaster* and D. busckii. One corresponding SB (gray box) is indicated with a gray connection between both species and corresponding BUSCOs are indicated with light red connections. The Hi-C profile of the corresponding SB is similar in both species.

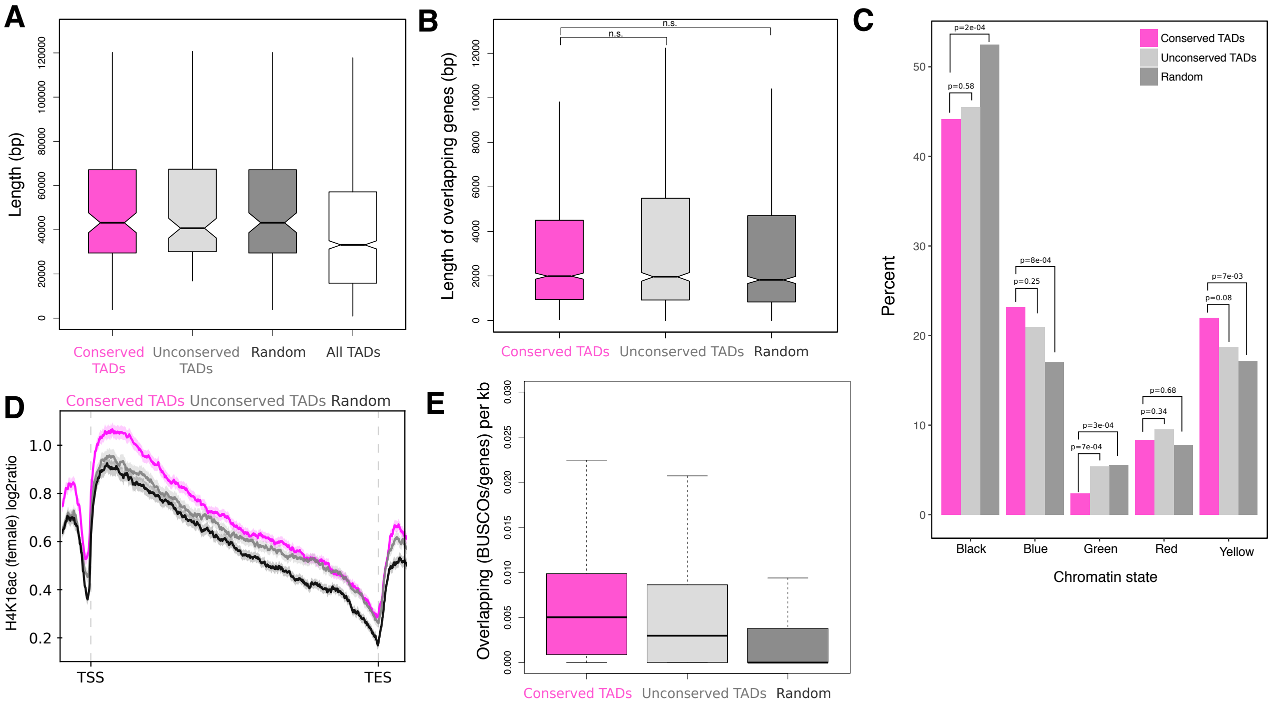

F

| *D. melanogaster* Motif | Logo | E-value | *D. virilis* Motif | Logo | E-value | *D. busckii* Motif | Logo | E-value |
|---|---|---|---|---|---|---|---|---|
| Ohler-5 | | 2.0e-37 | Ohler-5 | | 4.1e-46 | Ohler-5 | | 9.8e-43 |
| Beaf-32 | | 3.5e-32 | Beaf-32 | | 2.5e-42 | Beaf-32 | | 7.1e-35 |
| Ohler-1 | | 2.0e-30 | Ohler-8_dreme | | 1.4e-20 | ZIPIC | | 3.8e-32 |
| Ohler-8_dreme | | 1.5e-17 | GAATAGAAA | | 1.2e-14 | GAF | | 1.9e-29 |
| GAATAGAAA | | 8.3e-14 | ZIPIC | | 9.6e-14 | Ohler-8_dreme | | 2.9e-24 |
| GAF | | 3.2e-13 | Ohler-1 | | 1.1e-13 | Ohler-1 | | 5.5e-16 |
| Ohler-6 | | 1.2e-12 | Ohler-8 | | 8.3e-11 | Ohler-8 | | 1.1e-12 |
| ZIPIC | | 7.6e-11 | | | | CTCF | | 1.9e-12 |
| Ohler-8 | | 4.1e-09 | | | | | | |
| CTCF | | 8.3e-09 | | | | | | |

E-value < 1.0e-10

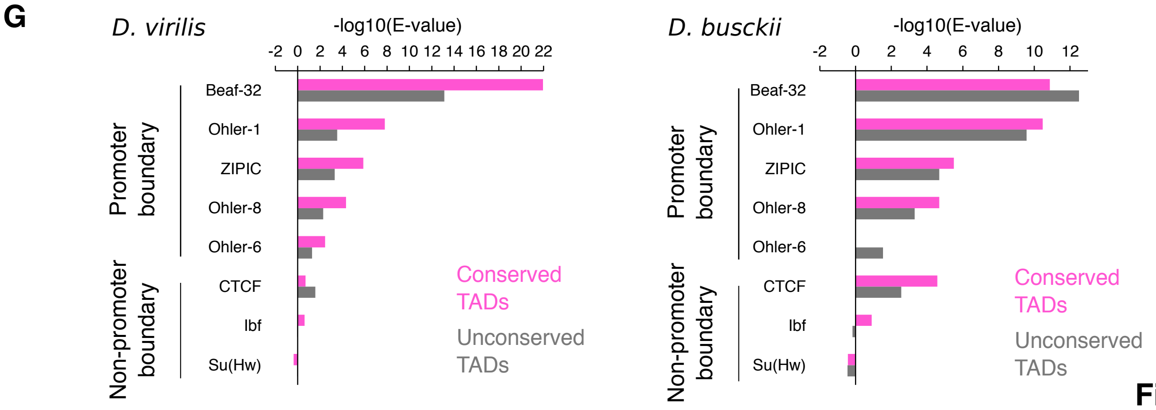

**Figure S6**

**Figure S6, related to Figure 4. Characterization of conserved TADs (pink) compared to unconserved TADs (grey) and random unconserved regions (dark grey).**

**(A)** Size distribution of the three test sets compared to all TADs (white).

**(B)** Length of genes overlapping with all test sets. This is a control for Figure 4B.

**(C)** Percentage of overlap of conserved TADs, unconserved TADs and random genomic regions with black, blue, green, red and yellow chromatin states defined in (Filion et al., 2010). *P*-values were calculated using Wilcoxon rank-sum tests on the chromatin states coverage (expressed in kb).

**(D)** Female H4K16ac signal over input of genes (transcription start site (TSS) to transcription end site (TES)) in conserved TADs (pink), unconserved TADs (grey) and random regions (black). ChIP-seq profiles show mean (thick line) and standard error (shadowed area) of input-normalized ChIP-seq enrichment along scaled genes and unscaled 1 kb before the TSS and after the TES.

**(E)** Fraction of BUSCOs in genes that overlap conserved TADs, unconserved TADs or random regions per kb.

**(F)** Enrichment of TAD boundary motifs in *D. melanogaster, D. virilis* and *D. busckii*. Enrichment analysis using AME from the MEME suite (McLeay and Bailey, 2010) for motif enrichment in 500 bp around TAD boundaries in *D. melanogaster, D. virilis* and *D. busckii*. We used motifs that were previously described in *D. melanogaster* (Ramírez et al., 2018) and analyzed their enrichment against shuffled input sequences as control, average odds score as the sequence scoring method and one-tailed Wilcoxon rank-sum test as the motif enrichment test.

**(G)** Analysis of motif enrichment of promoter and non-promoter boundary motifs at conserved TAD boundaries in *D. virilis* and *D. busckii* compared to unconserved TAD boundaries using AME from the MEME suite (McLeay and Bailey, 2010).

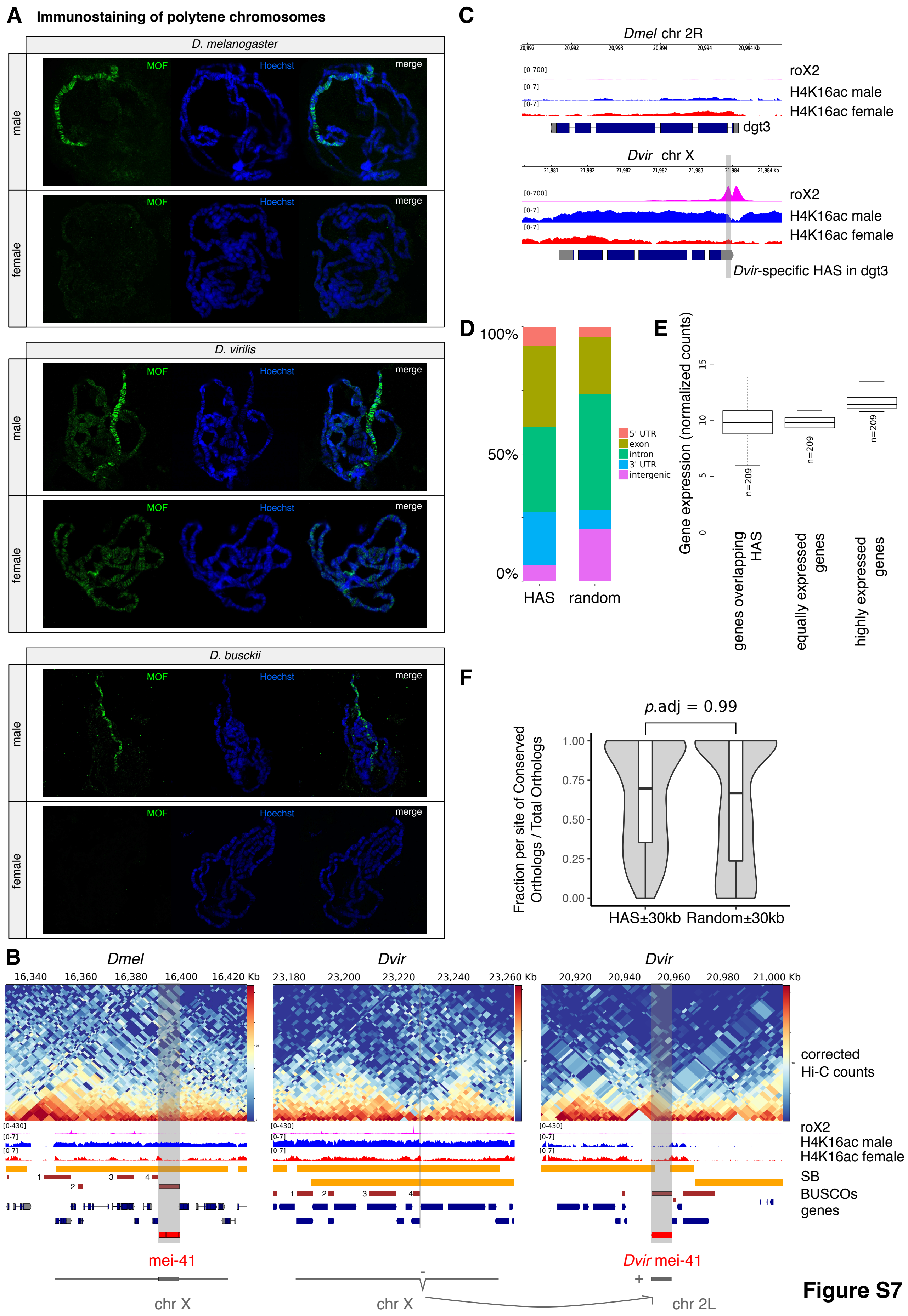
A Immunostaining of polytene chromosomes
D. melanogaster
D. virilis
D. busckii
male
female
MOF
Hoechst
merge
B
Dmel
Dvir
Dvir
16,340 16,360 16,380 16,400 16,420 Kb
23,180 23,200 23,220 23,240 23,260 Kb
20,920 20,940 20,960 20,980 21,000 Kb
corrected
Hi-C counts
roX2
H4K16ac male
H4K16ac female
SB
BUSCOs
genes
mei-41
Dvir mei-41
chr X
chr X
chr 2L
C
Dmel chr 2R
Dvir chr X
roX2
H4K16ac male
H4K16ac female
dgt3
Dvir-specific HAS in dgt3
D
100%
50%
0%
HAS
random
5' UTR
exon
intron
3' UTR
intergenic
E
Gene expression (normalized counts)
n=209
genes overlapping HAS
equally expressed genes
highly expressed genes
F
p.adj = 0.99
Fraction per site of Conserved Orthologs / Total Orthologs
1.00
0.75
0.50
0.25
0.00
HAS±30kb
Random±30kb

Figure S7

**Figure S7, related to Figure 5 and 6. Immunostaining of polytene chromosomes and HAS characterization.**

**(A)** Male and female polytene chromosomes were stained with MOF antibody (green) in *D. melanogaster, D. virilis* and *D. busckii*. DNA is counterstained with Hoechst (blue). MOF shows enrichment on the X chromosome compared to all other chromosomes in males but not females in all three species.

**(B)** Example gene that moved between the X chromosome and an autosome when comparing *D. melanogaster* and *D. virilis*. mei-41 is localized on chromosome X in *D. melanogaster* but on chromosome 2L in *D. virilis*. The surrounding genes on this SB on chromosome X maintained the same order (see corresponding BUSCOs numbered from 1-4 and gene track). mei-41 in *D. virilis* (Dvir GJ22445) is localized on chromosome 2L in between two surrounding SBs and only has a H4K16ac peak at the promoter (male and female) but the gene body is not covered with H4K16ac.

**(C)** The gene *dim gamma-tubulin 3* (Dmel\dgt3; Dvir\GJ18725) moved from chromosome 2L in *D. melanogaster* to the X chromosome in *D. virilis* and there gained roX2 binding in its 3' UTR. Thus, this HAS is a species-specific one.

**(D)** Fraction of HAS overlapping with 5' UTR, exon, intron 3' UTR or intergenic regions compared to random regions (n= 247).

**(E)** Gene expression (normalized counts) of genes overlapping with HAS compared to equally expressed genes and highly expressed genes (n = 209), referring to Figure 6E. Comparison of gene expression was performed using library size normalized RNA-seq counts from 14-20 h aged embryos from modENCODE datasets obtained from (Ramírez et al., 2018) and also available on the chorogenome web server (http://chorogenome.ie-freiburg.mpg.de/).

**(F)** Fraction of genes that are conserved compared to all genes per corresponding HAS pair between between *D. melanogaster* and *D. virilis* compared to random regions, both extended by 30 kb in 5' and 3' direction. Only genes with *D. melanogaster* to *D. virlis* orthologs were considered. The plotted distributions correspond to the fraction of orthologs in the *D. melanogaster* genome computed for each HAS or random site. The best matching pair of HAS or random region between both species was defined based on the highest (conserved orthologs / total orthologs) ratio. The displayed random regions boxplot and violin plot correspond to one of the 100 random sets analysed. The adjusted *p*-value is computed on all 100 random sets and was obtained by comparing the HAS distribution to every random set distribution using the Wilcoxon rank-sum test and Benjamini-Hochberg correction for multiple comparisons.

### Supplemental Tables

#### Table S1, related to Figure 1. Qualitative comparison of HiCAssembler with other Hi-C scaffolding tools.

| Hi-C scaffolding tool | LACHESIS | DNA Triangulation | GRAAL |
|---|---|---|---|
| Reference | 1 (Nov 2013) | 2 (Nov 2013) | 3 (Dec 2014) |
| Documentation | https://github.com/shendurelab/LACHESIS | https://github.com/NoamKaplan/dna-triangulation | https://github.com/koszullab/instaGRAAL |
| Aim (Proof-of-concept/Hi-C scaffolding tool) | Tool (not developed anymore) | Proof-of-concept | Tool (latest version instaGRAAL) |
| Requirements | unix environment | Python modules (numpy, scipy, scikit-learn) | NVIDIA graphic card |
| | gcc | | Cuda Toolkit |
| | zlib compression library | | PyCuda with OpenGL support |
| | boost C++ libraries | | Python dependencies (pip) |
| | SAMtools toolkit | | |
| Input | pre-assembled contigs or scaffolds | pre-assembled contigs or scaffolds of equally-sized bins | contigs or scaffolds split into bins of several restriction fragments |
| | aligned Hi-C data (.bam format) | Hi-C contact matrix | Hi-C contact matrix at several resolutions (pyramid file) |
| | ini file specifying several parameters | | |
| Predicts number of chromosomes | no | yes | yes |
| Orients contigs | yes | no | yes |
| Automatic misassembly detection | no | no | yes |
| Manual misassembly detection | no | no | no |
| Accounts for duplications | no | no | yes |
| Output | genome.fasta file | position of each bin (in an arbitrary units) | fasta file of genome assembly with the highest likelihood |
| | assembly metrics | | info about initial fragments |
| Visualization | no | no | assembly movie |
| Tested genome(s) | human, mouse and *Drosophila* | human | *Saccharomyces cerevisiae, Trichoderma reesei, human* |
| References | | | |
| 1 | https://www.nature.com/articles/nbt.2727 | | |
| 2 | https://www.nature.com/articles/nbt.2768 | | |
| 3 | https://www.nature.com/articles/ncomms6695 | | |
| 4 | https://bmcgenomics.biomedcentral.com/articles/10.1186/s12864-017-3879-z | | |
| 4 | https://doi.org/10.1101/261149 | | |
| 5 | http://science.sciencemag.org/content/356/6333/92 | | |

| Hi-C scaffolding tool | SALSA | 3D-DNA | HiCAssembler |
|---|---|---|---|
| Reference | 4 (Jan 2017) | 5 (Apr 2017) | this study |
| Documentation | https://github.com/machinegun/SALSA | https://github.com/theaidenlab/3d-dna | https://github.com/maxplanck-ie/HiCAssembler |
| Aim (Proof-of-concept/Hi-C scaffolding tool) | Tool (latest version SALSA2) | Tool | Tool |
| Requirements | Python 2.7 | Java version >=1.7 | Python dependencies (pip) |
| | BOOST libraries | Bash >=4 | |
| | networkx | GNU Awk >=4.0.2 | |
| | | GNU coreutils sort >=8.11 | |
| | | LastZ version 1.03.73 (for diploid mode only) | |
| | | Python >=2.7 (for chromosome number-aware splitter module only) | |
| | | scipy numpy matplotlib (for chromosome number-aware splitter module only) | |
| Input | pre-assembled contigs or scaffolds | pre-assembled contigs or scaffolds | pre-assembled contigs or scaffolds |
| | aligned Hi-C data (.bed format) | Hi-C interaction matrix in .hic format (from Juicebox Hi-C analysis software) | Hi-C interaction matrix in .h5 format (from HiCExplorer Hi-C analysis software) |
| Predicts number of chromosomes | yes | yes | yes |
| Orients contigs | yes | yes | yes |
| Automatic misassembly detection | yes | yes | yes |
| Manual misassembly detection | no | yes | yes |
| Accounts for duplications | no | under development | no |
| Output | genome.fasta file | genome.fasta file | genome.fasta file |
| | | .hic file | .h5 file |
| | | scaffold boundary files | liftover.chain file (allows liftover of data from inital contigs to assembly) |
| | | tracking file of modifications to the input contigs (.assembly file) | files for visualization of assembly path graph (.graphml files) |
| Visualization | no | Juicebox Assembly Tools (JBAT) can be used for visualization | automatic visualization at each assembly iteration |
| Tested genome(s) | human, goat | *Aedes aegypti, Culex quinquefasciatus* | *Drosophila busckii, Drosophila virilis* |
| References | | | |
| 1 | | | |
| 2 | | | |
| 3 | | | |
| 4 | | | |
| 4 | | | |
| 5 | | | |

**Table S2, related to Figure 1. Sequenced and filtered valid reads of Hi-C samples.** Upper panel contains numbers of sequenced reads for each sample. Some samples were sequenced in two runs to get higher sequencing depth which is indicated as 'Run_1' and 'Run_2'. Lower panel contains valid used reads for each sample after creation of the Hi-C contact matrix using `hicBuildMatrix` from HiCExplorer (see methods: Creation of Hi-C contact matrix or (Ramírez et al., 2018)), as well reported in percent with respect to pairs sequenced.

| | Sample | *Dmel* Rep1 | *Dmel* Rep2 | *Dvir* Rep1 | *Dvir* Rep2 | *Dbus* Rep1 | *Dbus* Rep1 | *Dbus* 21-23h |
|---|---|---|---|---|---|---|---|---|
| Sequenced reads | Run 1 | 55,222,595 | 68,801,187 | 64,177,938 | 60,426,728 | 6,491,637 | 22,258,838 | 69,290,579 |
| | Run 2 | 85,848,394 | 52,938,683 | 0 | 0 | 107,801,339 | 77,256,684 | 0 |
| | Total | 141,070,989 | 121,739,870 | 64,177,938 | 60,426,728 | 114,292,976 | 99,515,522 | 69,290,579 |
| | Total Rep1&2 | 262,810,859 | | 124,604,666 | | 213,808,498 | | 69,290,579 |

| | Sample | *Dmel* Rep1 | *Dmel* Rep2 | *Dvir* Rep1 | *Dvir* Rep2 | *Dbus* Rep1 | *Dbus* Rep1 | *Dbus* 21-23h |
|---|---|---|---|---|---|---|---|---|
| Useful reads HiCExplorer 1.8.1 | Run 1 | 9,349,695 | 12,242,090 | 26,929,877 | 24,857,426 | 2,216,509 | 5,687,685 | 26,811,828 |
| | Run 2 | 14,611,364 | 9,501,536 | 0 | 0 | 37,669,762 | 18,934,139 | 0 |
| | Total | 23,961,059 | 21,743,626 | | 51,787,303 | 39,886,271 | 24,621,824 | 26,811,828 |
| | Total Rep1&2 | 45,704,685 | | 51,787,303 | | 64,508,095 | | 26,811,828 |

| | Sample | *Dmel* Rep1 | *Dmel* Rep2 | *Dvir* Rep1 | *Dvir* Rep2 | *Dbus* Rep1 | *Dbus* Rep1 | *Dbus* 21-23h |
|---|---|---|---|---|---|---|---|---|
| Useful / Sequenced reads | Run 1 (%) | 16.9 | 17.8 | 42.0 | 41.1 | 34.1 | 25.6 | 38.7 |
| | Run 2 (%) | 17.0 | 18.0 | NA | NA | 34.9 | 24.5 | NA |
| | Average (%) | 17.4 | | 41.6 | | 29.8 | | 38.7 |

**Table S3, related to Figure 3A. Fischer two-tail test results for the overlap of TADs with SBs.**

| Comparison | Species | Condition | Fisher two tail *p*-value | Overlapping proportion |
|---|---|---|---|---|
| *D. busckii* vs *D. melanogaster* | *D. busckii* | Called TADs | 2.29E-62 | 0.17 |
| | | Shuffled SBs | 5.55E-01 | 0.07 |
| | | Shuffled TADs | 6.76E-01 | 0.07 |
| | | Shuffled TADs & SBs | 3.00E-01 | 0.07 |
| | *D. melanogaster* | Called TADs | 4.84E-193 | 0.22 |
| | | Shuffled SBs | 5.88E-01 | 0.06 |
| | | Shuffled TADs | 5.86E-02 | 0.07 |
| | | Shuffled TADs & SBs | 9.70E-01 | 0.06 |
| *D. busckii* vs *D. melanogaster* | *D. virilis* | Called TADs | 1.41E-65 | 0.13 |
| | | Shuffled SBs | 5.85E-02 | 0.06 |
| | | Shuffled TADs | 6.81E-01 | 0.05 |
| | | Shuffled TADs & SBs | 7.71E-01 | 0.05 |
| | *D. melanogaster* | Called TADs | 6.33E-212 | 0.22 |
| | | Shuffled SBs | 8.36E-01 | 0.06 |
| | | Shuffled TADs | 5.04E-01 | 0.06 |
| | | Shuffled TADs & SBs | 7.75E-01 | 0.06 |
